# The feedback loop of the *transformer* gene results in proper feminization under multiple sex determination signals in the ant, *Vollenhovia emeryi*

**DOI:** 10.1101/2022.12.14.520459

**Authors:** Misato Okamoto Miyakawa, Hitoshi Miyakawa

**Affiliations:** Center for Bioscience Research and Education, Utsunomiya University, 350, Minemachi, Utsunomiya, Tochigi 321-8505, Japan

**Keywords:** Complementary sex determiner, haplodiploidy, *ml*-CSD, *transformer*, hymenoptera

## Abstract

Organisms that reproduce sexually have evolved highly sophisticated mechanisms to generate two sexes. Some hymenopterans (such as ants, bees, and wasps) have a complementary sex-determination system in which heterozygosity at one CSD locus induces female development, whereas hemi- or homozygosity at the locus induces male development. This system can generate a high cost of inbreeding, as individuals that are homozygous at the locus become sterile, diploid males. On the other hand, some hymenopterans have evolved a multi-locus, complementary, sex-determination system in which heterozygosity in at least one CSD locus induces female development. This system effectively reduces the proportion of sterile diploid males; however, how multiple primary signals pass through a molecular cascade to regulate the terminal sex-determination gene has remained unclear. To clarify this matter, we investigated the molecular cascade in the ant, *Vollenhovia emeryi*, with two CSD loci. Here we show that *transformer* (*tra*) is necessary for proper feminization. Expression and functional analysis showed that individuals heterozygous in at least one of the two CSD loci develop into females. Individuals heterozygous at only a single CSD locus do not develop into sexual intermediates, because the signal derived from the heterozygous CSD is amplified by a positive-feedback splicing loop of *tra*. Our data also demonstrate that *tra* controls splicing of *doublesex* (*dsx*), which is involved in sexual differentiation. We suggest a cascade model to arrive at a binary determination of sex under multiple primary signals.

## Introduction

Deciphering the means by which molecular mechanisms of sex determination have diversified is a fundamental problem of biology. While primary sex determination signals are diverse and can evolve rapidly [1, 2], basic structures of sex determining programs are somewhat similar across species [3, 4]. In organisms that reproduce sexually, genes involved in sex determination are fundamentally bipotent, having the potential to develop into either sex, depending on primary signals [5]. Accordingly, the sex determination process requires sophisticated gene regulatory networks to ensure proper development into one sex or the other [5, 6].

Haplodiploidy, is one of the major sex determination systems that evolved independently in several animal groups, including once at the origin of the insect order, Hymenoptera [7]. One of the molecular mechanisms underlying haplodiploidy is complementary sex determination (CSD) in which individuals heterozygous at the CSD locus become females, while those that are homo- or hemizygous become males [7–9]. Although molecular characterization of the CSD locus has been accomplished only in the honeybee, *Apis mellifera* [10–12], involvement of the CSD locus in sex determination is also predicted in other hymenopterans since inbreeding crosses result in diploid males [13, 14]. The haplodiploid sex determination system allows females to control the sex ratio [15, 16]; however, involvement of the CSD locus imposes a high inbreeding cost (up to 50%; *Figure 1*A), since diploid males resulting from homozygosity at the CSD locus are typically sterile or inviable, and small populations can fall into an extinction vortex [8,17–20]. On the other hand, some hymenopterans (ants and wasps) have evolved a CSD-independent sex-determination system [21–25] or multi-locus CSD (*ml*-CSD) systems [26–31]. As a result, diploid male production is avoided or reduced under inbreeding. In *ml*-CSD, diploid individuals that are heterozygous for at least one of multiple CSD loci develop into females, whereas individuals that are haploid hemizygous or diploid homozygous at all loci develop into males [26–31]. For example, in the ant *Vollenhovia emeryi*, with two CSD loci, diploid males in inbred crosses are suppressed to 25% (*Figure 1*B)[28]. In another example, in the parasitoid wasp, *Lysiphlebus fabarum*, with four CSD loci, diploid males in inbred crosses are theoretically suppressed to 6.25% [30]. Importantly, sexual intermediates never arise, even when feminization (heterozygous CSD) and masculinization (homozygous CSD) primary signals occur simultaneously [28, 30]. Although empirical and theoretical studies have shown that *ml*-CSD systems suppress the cost of inbreeding by reducing the chance that all CSD loci become homozygous [18, 32], molecular sex-determination mechanisms controlled by multiple primary signals derived from CSD loci are still unknown.

**Figure 1.**
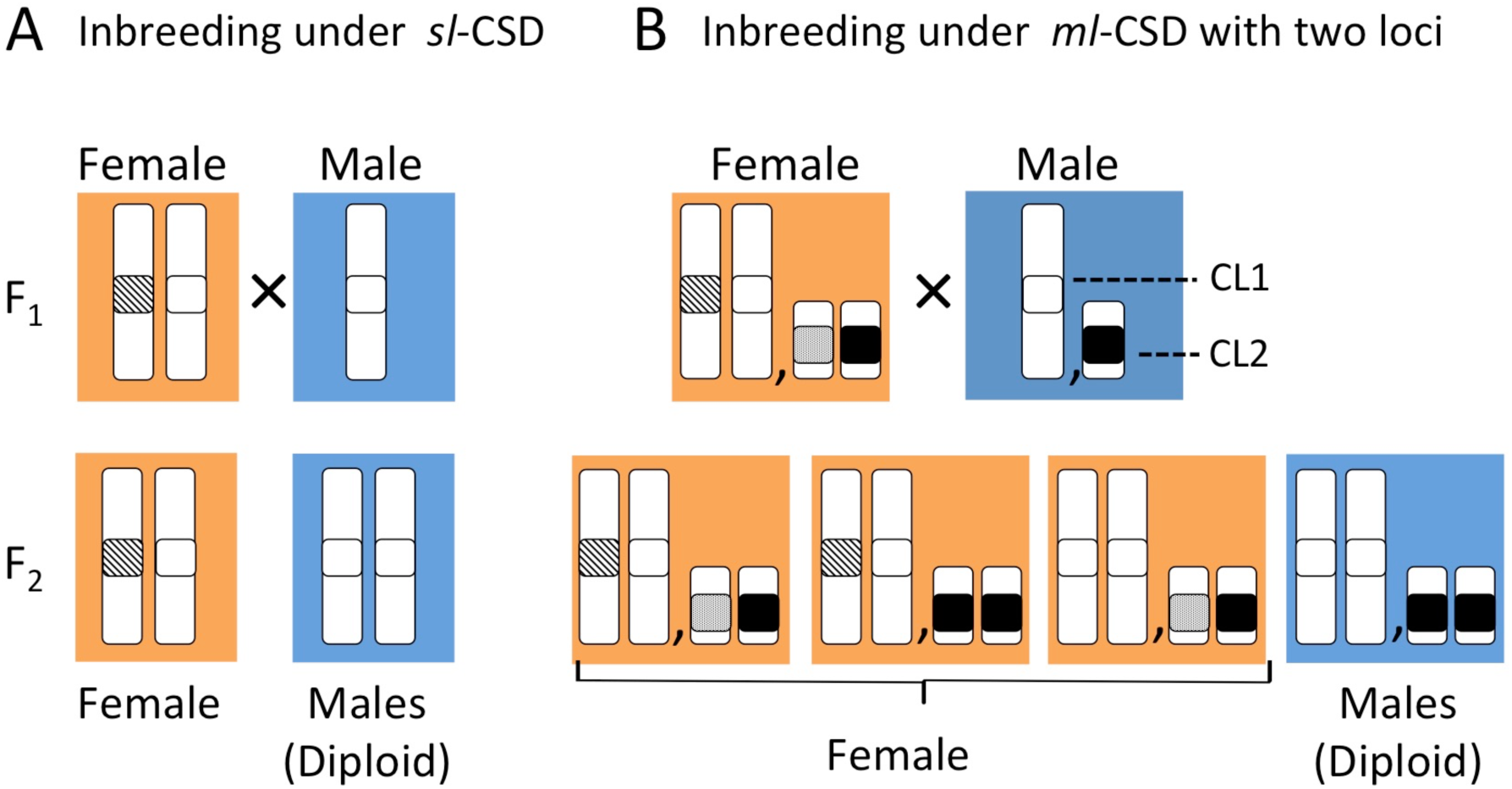
Allele patterns of offspring produced by inbreeding crosses. (A) Under single-locus CSD (*sl*-CSD), 50% of all offspring became sterile, diploid males, homozygous at a CSD locus. (B) Under multi-locus CSD (*ml*-CSD) with two loci, the inbreeding load is half of the *sl*-CSD system, since production of diploid males homozygous at two loci is limited to 25% of all offspring.

*Vollenhovia emeryi* is particularly well suited to investigations of molecular sex determination because of its unusual reproductive system, in addition to relative ease of rearing and inbreeding (*Figure 2*) [33–35]. Queens are produced parthenogenetically and are clones of their mothers (*Figure 2*a,c), while males are produced androgenetically and are genetically identical to their fathers (*Figure 2*b,e). On the other hand, workers and some queens are produced sexually (*Figure 2*d)[33,34,36]. Our previous genomic study suggested that primary sex determination signals are different between such sexually produced females and clonal queens (*Figure 2*)[28]. Sex determination in sexually produced females in *V. emeryi* relies on two CSD loci [28] and is therefore an excellent model for investigating the molecular sex determination system controlled by *ml*-CSD [37]. We identified two quantitative trait loci (QTL), which explained the phenotypic variance in diploid males obtained from inbreeding crosses using sexually produced queens (*Figure 2*d) with their brothers (*Figure 2*e). We also demonstrated that only a quarter of all inbred offspring became diploid males as mentioned above [28]. One of the two QTLs (hereafter, CL1) contains two tandem *transformer* (*tra*) homologs, as in honeybees and other ants [28,38,39]. The other QTL (hereafter, CL2) does not contain these orthologs, implying that molecular substances of CL2 have newly acquired CSD-like functions perhaps by convergence [28]. We also showed that splicing of *doublesex* (*dsx*) is responsible for sex-specific traits in *V. emeryi*, suggesting that two independent primary signals on CL1 and CL2 are finally integrated into *dsx* [37]. On the other hand, female determination system in clonal queens (*Figure 2*a,c) could not be clearly explained by state of two CSD loci, and the details of the primary sex-determining signals are unknown [28].

**Figure 2.**
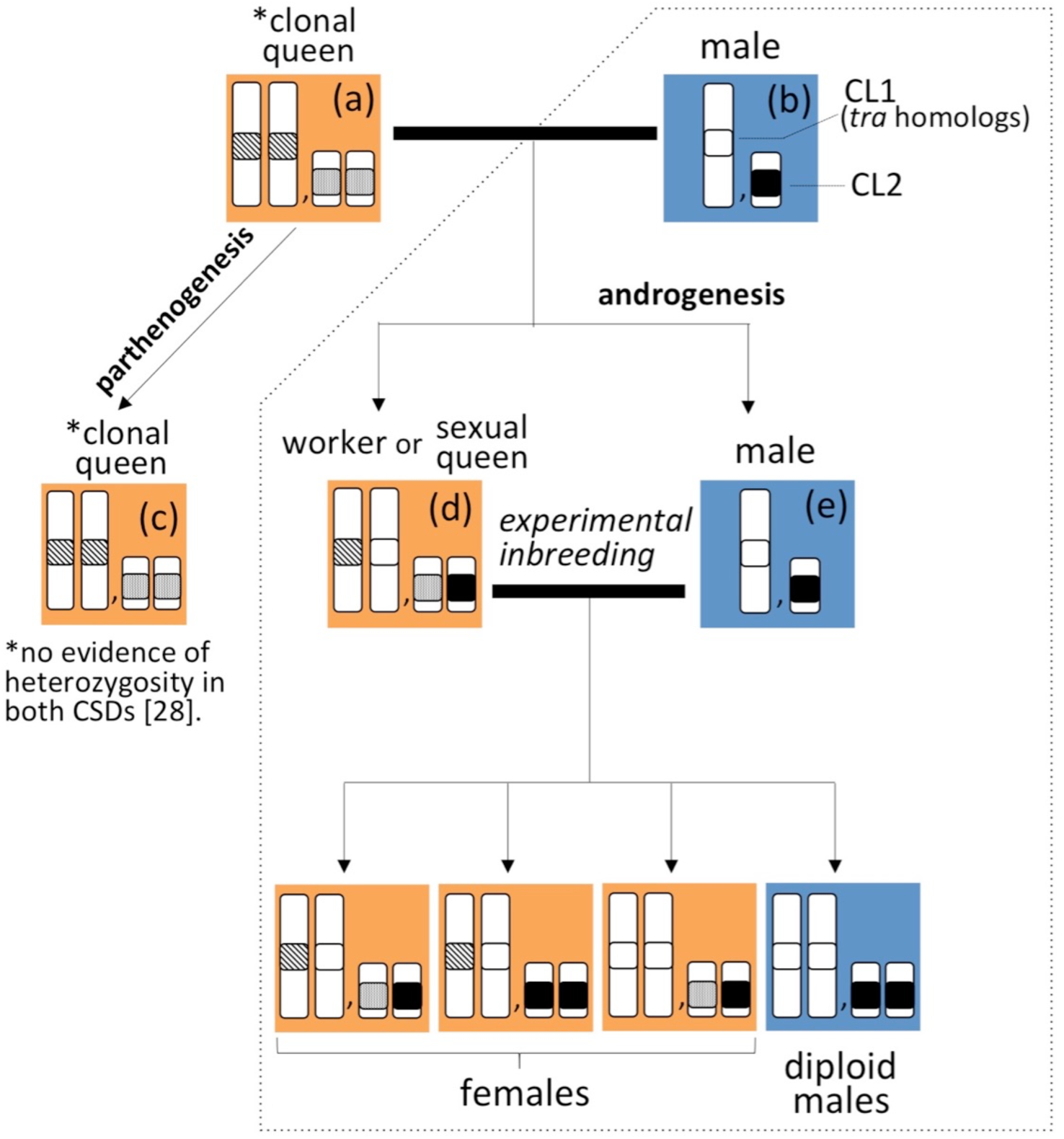
Extraordinary reproduction system and design for experimental crosses in *V. emeryi*. *V. emeryi* has an unusual reproduction system with queen parthenogenesis and male androgenesis. Crosses of parental queens (a) and males (b) produce three types of offspring: parthenogenetically produced clonal queens (c) with only maternal genomes, workers and sexual queens (d) with parental genomes, and androgenetically produced males (e) with only paternal genomes. To obtain new offspring for gene expression analysis, we experimentally crossed sexual queens (d) and androgenetic males (e), which is equivalent to a backcross (inbreeding). Previous QTL mapping and diploid male ratio support a two-locus model of sex determination *in V. emeryi* (encircled by a dashed line). However, homozygous of *tra* homologs at CL1 and homozygous of exonic region at CL2 suggest separate sex determination mechanism in only clonal queens [28]. In this study, we also crossed clonal queens and androgenetic males, which is equivalent to outbreeding. We refer to this outbreeding as “genetic-outbreeding” to distinguish it from crossing using geographically separated males and queens in previous study [28]. Although the cytological mechanisms are unknown, clonal queen eggs are predicted to be produced by fusion of two of the four meiotic products, while clonal male eggs are predicted to be generated by elimination of maternal genomes from inseminated eggs or insemination of enucleated eggs and sperm [63].

In this study, we have challenged to discover molecular mechanisms enabling *ml*-CSD system in *V. emeryi* using offspring produced by inbreeding crosses. We predicted that there would be a molecular intermediate that converts two upstream signals into a binary response and directs the splicing of *dsx*. For example, in holometabolous insects, *tra* itself is alternatively spliced so that it produces an active protein only in females, whereas in males, a premature stop codon results in a truncated, inactive Tra protein [40–44]. Thus, the female *dsx* isoform requires active intervention by *tra*, whereas the male-specific *dsx* isoform is produced by default. Notably, active Tra protein has an additional positive feedback activity that directs processing of female-type mRNAs, suggesting that they function not only in sex determination, but also in maintaining the determined sex throughout development [45, 46]. In honeybees, the *tra* gene is duplicated into two paralogs called *feminizer* (*fem*) and *complementary sex determiner* (*csd*). The former has the conserved *tra* function, and in contrast, the latter has acquired a novel function in sex determination. In *A. mellifera*, *csd* derived from heterozygous CSD regulates the female type of *fem* splicing followed by the female type of *dsx* splicing, while *csd* derived from hemi/homozygous CSD regulates the male type of *fem* splicing followed by the male type of *dsx* splicing [12, 47]. Functional analysis showed that the female sex determination pathway is exclusively induced by *csd* and maintained via a positive-feedback loop of *fem* [10]. Accordingly, we hypothesize that in *V. emeryi*, diploid individuals employing homo- and heterozygous CSDs develop into females, since the primary signal derived from only a heterozygous CSD is amplified by positive feedback at the level of *tra*.

Here, we identified and investigated the function of *tra* in the sex determination cascade in *V. emeryi.* In this study, we first obtained the full sequences of the *tra* homolog and confirmed sex-specific gene expression. We next conducted expression analysis for each individual produced by inbreeding crosses to compare allele patterns of the two CSD loci (CL1 and CL2), splicing variants of *tra* and *dsx*, and sexual phenotypes. We found that expression of *tra* and *dsx* in females homozygous at one of the two CSD loci does not differ from that of females heterozygous at both CSD loci. Introduction of the female-type isoform of *tra* mRNAs in males induces splicing of endogenous *tra* pre-mRNAs into the female-type isoform. We also showed that inhibition of *tra* function caused sex reversal from female to male. Based on these results, we suggest that *tra* has the capacity to transduce the initial signal derived from the heterozygous CSD locus by a positive-regulatory-feedback loop and is the key element to realize proper feminization under multiple primary sex determination signals in *V. emeryi*.

## Results

### Characterization of *transformer* homologs in *V. emeryi*

We cloned full-length cDNA sequences corresponding to *V. emeryi* gene models of *traA* (*NCBI NM_001310035.1* [28]) and *traB* (NCBI *NM_001310036.1* [28]) (*Figure 3A*). Of the two homologs, only *traB* shows sex-specific splicing, as in *tra* in other holometabolous insects [12,45,48]. These data suggested that *traB* is an ortholog of *tra* and that *traA* is a paralog of *traB*. Thus, we regarded *traB* as *tra* and here *traA* without sex-specific splicing is termed a “*tra-like*” gene. Amplified fragments in the *tra-like* RT-PCR experiments were comprised of 9 exons, and the exon-intron structure was identical between sexes. In *tra-like* isoforms, the sex-determination protein N-terminal domain (pfam12278) [11] and the complementary sex-determiner protein region domain (pfam11671)[11, 49] were located from exon 2 to 3, and from exon 5 to 9 (*Figure 3A*). In addition, a coiled-coil motif was predicted in exon 2 by InterProScan (https://www.ebi.ac.uk/interpro/search/sequence/) (*Figure 3A - Supplementary figure 1*). Although exon-intron structure of *tra-like* is identical between sexes, lengths of mRNAs were different (females: 1949 nt encoding 477 aa CDS, males: 1666 nt encoding 467 aa CDS). Translated amino acid sequences of *tra-like* showed different numbers of repeated amino acids (VSN, HQN, etc.) at the end of the complementary sex-determiner protein region domain in exon 9 (*Supplementary figure 1A*). Using samples from three geographically separated sites, we conducted additional RT-PCR to amplify the repeated region of *tra-like* and detected four types of sequences with different numbers of amino acid repeats (*Supplementary figure 1B*). At all collection localities, clonal queens and males mating naturally in the field possessed different types of *tra-like*. The domains and motifs in *tra-like* are similar to those of honeybee *csd*. On the other hand, female and male *tra* mRNAs were comprised of 11 exons (2246 nt encoding 452 aa CDS) and more than 14 exons (>2027 nt encoding 191 aa CDS), respectively (*Figure 3*A). In the female isoform, the sex-determination protein N-terminal domain (pfam12278) [11] and the complementary sex-determiner protein region domain (pfam11671) [11, 49] were located from exon 3 to 4, and in exon 9 (*Figure 3A*). In all detected *tra* male isoforms, the termination codon arose at exon 4 or 5, and male-specific exons occurred in the region corresponding to the intron in the female isoform (*Figure 3A*-*Supplementary figure 2*). Amplifications of multiple *tra*^M^ isoforms shown in several males suggest that males synthesize multiple *tra*^M^ isoforms simultaneously (*Supplementary figure 2*). This pattern, an earlier termination codon in the male isoform, resulting in production of a full-length protein in females, but not in males, follows the same pattern as *tra* in other holometabolous insects [12,45,48].

**Figure 3.**
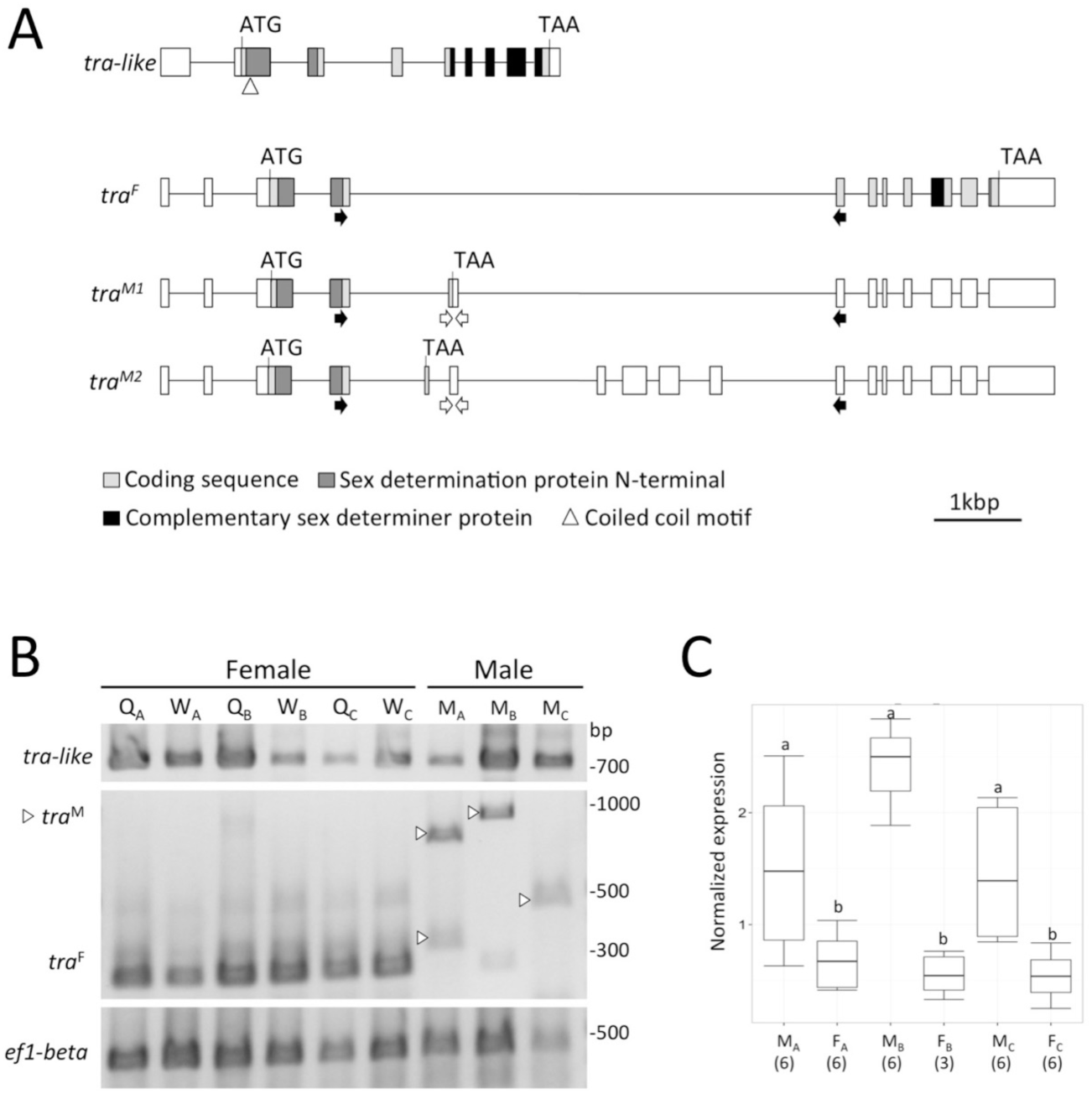
Genomic organization and expression of two *tra* homologs, *tra* and *tra-like* in *V. emeryi.* (A) Exon (boxes) and intron (lines) structure diagram of *tra-like* and *tra*. Only *tra* shows female- (*tra*^F^) and male- (*tra*^M^) specific isoforms. Other male isoforms of *tra* are shown in *Supplementary figure 2*. Exon and intron structure of *tra-like* is identical between sexes, but the length of the last exon is different. The translated amino acid sequence of *tra-like* showed a different number of repeated motifs at the last exon (*Supplementary figure 1*). A coiled-coil motif was predicted in *tra-like*, but not in *tra*. Black and white arrows indicate primer regions used in RT-PCR (B) and qPCR (C), respectively. (B) Expression of female (queen Q_N_/worker W_N_) and male (M_N_) splice forms of *tra-like* and *tra* by RT-PCR. Subscripts a - c indicate the site of collection (a: Ishikawa Prefecture, b: Tokyo Prefecture, c: Tochigi Prefecture). (C) Normalized gene expression (median ± SD) of *tra*^M^ inferred by qPCR. Adult female (F_N_) and male (M_N_) from each site of collection as with (B). Different letters indicate significant differences among groups (Tukey-Kramer test, P < 0.05). Sample sizes are given in parentheses. Boxplots show the median and interquartile ranges (IQR) and 1.5 IQR. The following figure supplements are available for Figure 3: **Supplementary figure 1.** cDNA and amino acid sequence of *tra-like* in *V. emeryi*. **Supplementary figure 2.** Exon (boxes) and intron (lines) structure diagram of *tra* in *V. emeryi*.

To confirm whether these *tra* isoforms detected in *V. emeryi* are expressed in a sex-specific manner, we performed an RT-PCR comparison between male and female isoforms with a primer pair designed to amplify sex-specific exons. Based on the gene structure, amplification of fragments of 274 bp and more than 326 bp was expected in females and males (*Figure 3B*-*source data 1*). A fragment <300 bp was amplified in all female samples, while a fragment >300 bp was predominantly amplified in all male samples (*Figure 3B-source data 1*). These results suggest that the obtained *tra* sequences were indeed isoforms involved in sexual differentiation, and are hereafter referred to as female and male isoforms, *tra*^F^ and *tra*^M^. On the other hand, in RT-PCR of *tra-like* with a primer pair designed to amplify all exons within the ORF, amplicons of 734 bp and 738 bp were expected in females and males. A fragment of approximately 700 bp was amplified in all females and males.

Next, we analyzed expression profiles of *tra* isoforms more precisely using qPCR. *tra*^M^ is expressed more highly and significantly in males than in females. (P < 0.05; *Figure 3C*). *tra*^F^ expression could not be detected by qPCR because specific primers could not be designed for the female-specific region. These data suggest that *V. emeryi tra* shows sex-specific splicing variants and expression, as with *dsx* [37].

### Gene expression, allele types of both CSD loci, and phenotypes

The relation between allele types of two CSD loci (CL1 and CL2) and expression patterns of *tra* and *dsx* were tested using offspring produced by inbreeding (*Figure 1B*, *Figure 2*). First, we established the high-resolution melting curve (HRM) analysis method for zygosity testing of two CSD loci. Locations of SNPs for the HRM analysis, which were screened from previous QTL data [28], are shown in *Figure 4*. The amplicon of CL1 (116 bp) contains two SNPs that are at the 126,516th and the126,539th nucleotides of *NCBI NW_011967057.1* (*Figure 4*). Combinations of these SNPs were the following two cases: (1) “T” at the 126,516th and “C” at the126,539th (written as “T..C”), (2) “C” at the 126,516th and “T” at the126,539th (“C..T”). The amplicon of CL2 (111 bp) contains one SNP at the 95,390th nucleotide of *NCBI NW_011967112.1* (*Figure 4*). Queens used for inbreeding crosses were heterozygous T..C/C..T and T/C at CL1 and CL2, respectively. On the other hand, males used for inbreeding crosses (androgenetic male in *Figure 2*) were hemizygous T..C and C at CL1 and CL2, respectively. Females produced by inbreeding crosses were heterozygous T..C/C..T or homozygous T..C/T..C at CL1, and heterozygous T/C or homozygous C/C at CL2 (*Figure 5*). All males produced by inbreeding crosses were only homozygous T..C/T..C and C/C at CL1 and CL2, respectively (*Figure 5*). To confirm the relationship between allele type of two CSD loci predicted by HRM analysis and ploidy level of offspring produced by inbreeding crosses, we measured relative quantities of DNA in individuals with each allele type. Flow cytometry showed distinct peaks between inbred offspring and androgenetic males, which are predicted diploid (P2) and haploid (P1) (*Supplementary figure 3*). Accordingly, males produced by inbreeding crosses are diploid, and are homozygous at both CSD loci, CL1 and CL2.

**Figure 4.**
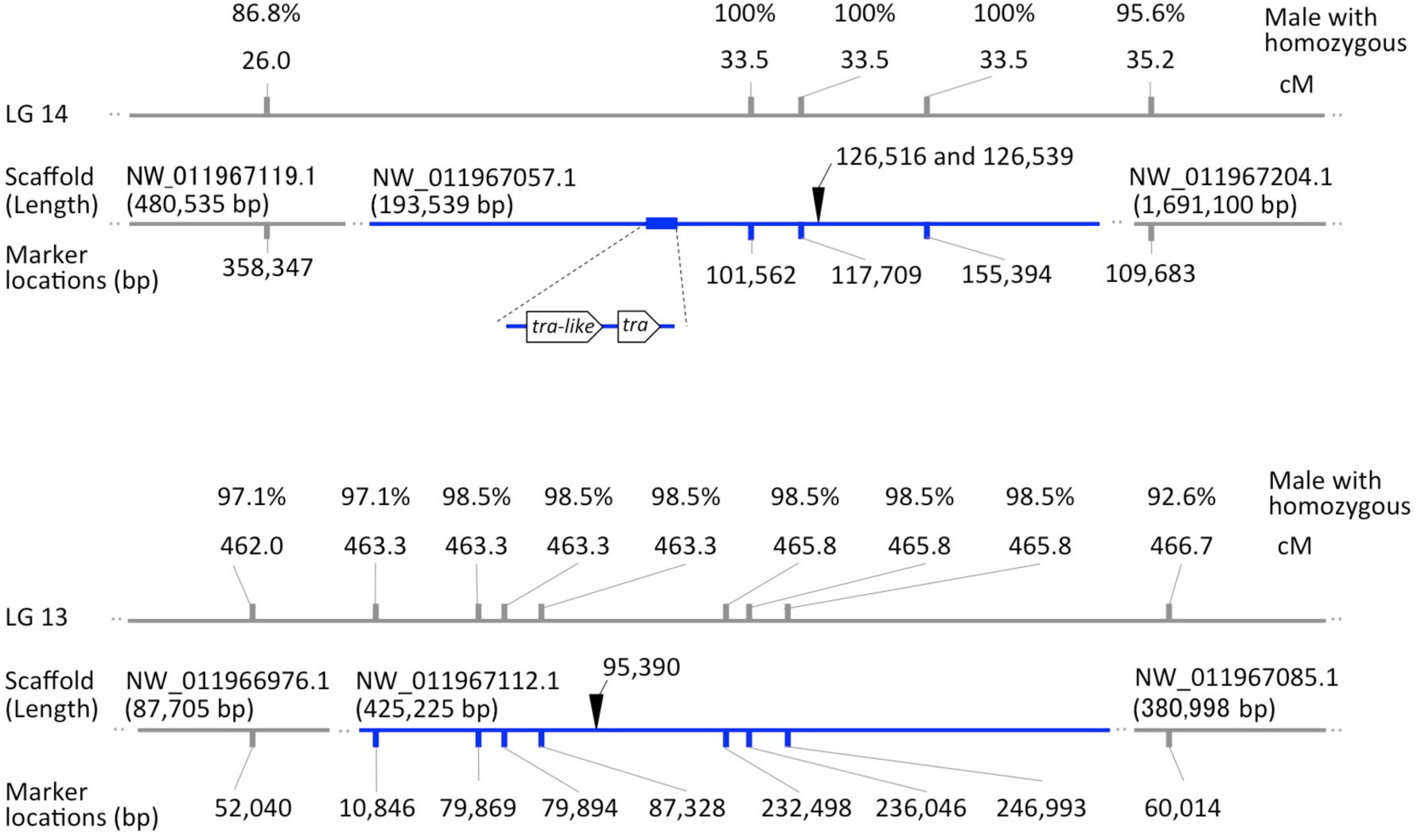
Location of SNPs for HRM analysis within QTLs associated with sex determination genes. Blue lines show two QTLs, CL1 and CL2, located in separate linkage groups, LG 14 and LG 13 [28]. Phenotypes of diploid males (n=68) were strongly associated with homozygosity at these loci. SNPs used for HRM analysis are indicated by black arrowheads. These SNPs on CL1 are located within three tightly linked markers [0 centimorgans (cM) apart] associated with QTL and located 42kb from *tra-like*. SNPs for HRM analysis on CL2 are located within seven markers (3.2 cM) associated with another QTL. In CL1, *tra-like* and *tra* are encoded from 79,750bp to 84,722bp and from 87,411bp to 97,941bp, respectively.

**Figure 5.**
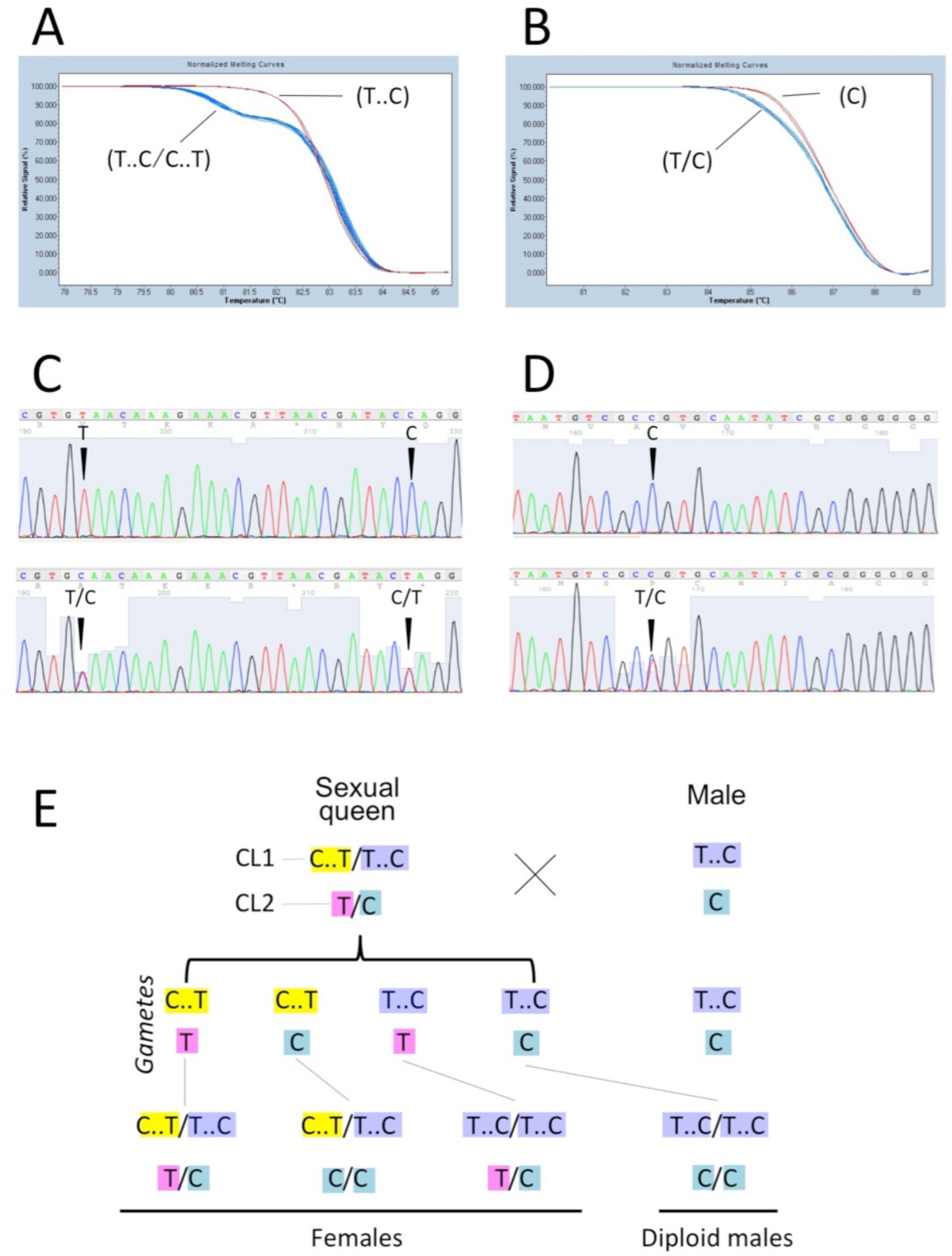
Genotyping by HRM analysis using offspring produced by inbreeding. Normalized melting curves of amplicons containing SNPs of CL1 (A) and CL2 (B). Red and blue curves are homo- and heterozygous at the SNPs, respectively. Raw sequencing view of SNP regions at CL1 (C) and CL2 (D). Graphical summary of SNPs of individuals obtained from this experiment (E).

Next, we investigated frequencies of individuals with each allele type of two CSD loci using inbred offspring derived from three geographically separated sites. Females were heterozygous at one or both CSD screening regions, with the exception of a few specimens from Ishikawa Prefecture, in which two of the 139 females produced by inbreeding proved homozygous at both screening regions, as with diploid males. These exceptions may result from occurrence of recombination in the screening SNPs or due to malfunction of the downstream sex differentiation cascade. Proportions of females with each allele type (CL1/CL2: hetero/hetero, hetero/homo, homo/hetero) were equal (Ishikawa: χ^2^(2) = 1.503, p = 0.472, Tochigi: χ^2^(2) = 1.902, p = 0.386, Tokyo: χ^2^(2) = 1.714, p = 0.424; *Supplementary figure 4*). All males on which we performed HRM analysis proved homozygous at both CSD screening regions. The results of HRM analysis suggest that both CSD screening regions are strongly linked to primary sex-determination genes and that the method is applicable to zygosity testing of the two CSD loci in *V. emeryi*.

Finally, we tested the relationship between the allele types of two CSD loci and the expression pattern of *tra* and *dsx*. In *tra*^F^ expression, RT-PCR performed on male and female isoforms showed female-specific isoform amplification in all female samples, each with three alleles types at CSD loci (CL1: hetero / CL2: hetero, CL1: hetero / CL2: homo, CL1: homo / CL2: hetero, *Figure 6A*-*source data 2*, *Supplementary figure 5- source data 4*). As a result of qPCR, irrespective of alleles at CSD loci, *dsx*^F^ was highly expressed in all female samples (*Figure 6B*). On the other hand, *tra*^M^ and *dsx*^M^ were highly expressed in all male samples (*Figure 6B*), which comprised haploid androgenetic males with (CL1: hemi / CL2: hemi) and diploid males with (CL1: homo / CL2: homo). These data revealed that even though *V. emeryi* females show three allelic combinations at the two primary sex-determination loci (*Figure 1B*), sex-specific splicing and expression of *tra* and *dsx* correspond exactly to the appearance of sexual individuals. In other words, *tra*^F^ and *dsx*^F^ expression levels in females that are homozygous at one of the two CSD loci do not differ from females that are heterozygous at both CSD loci. Similarly, neither expression nor splicing of *tra* and *dsx* differed between diploid and haploid males.

**Figure 6.**
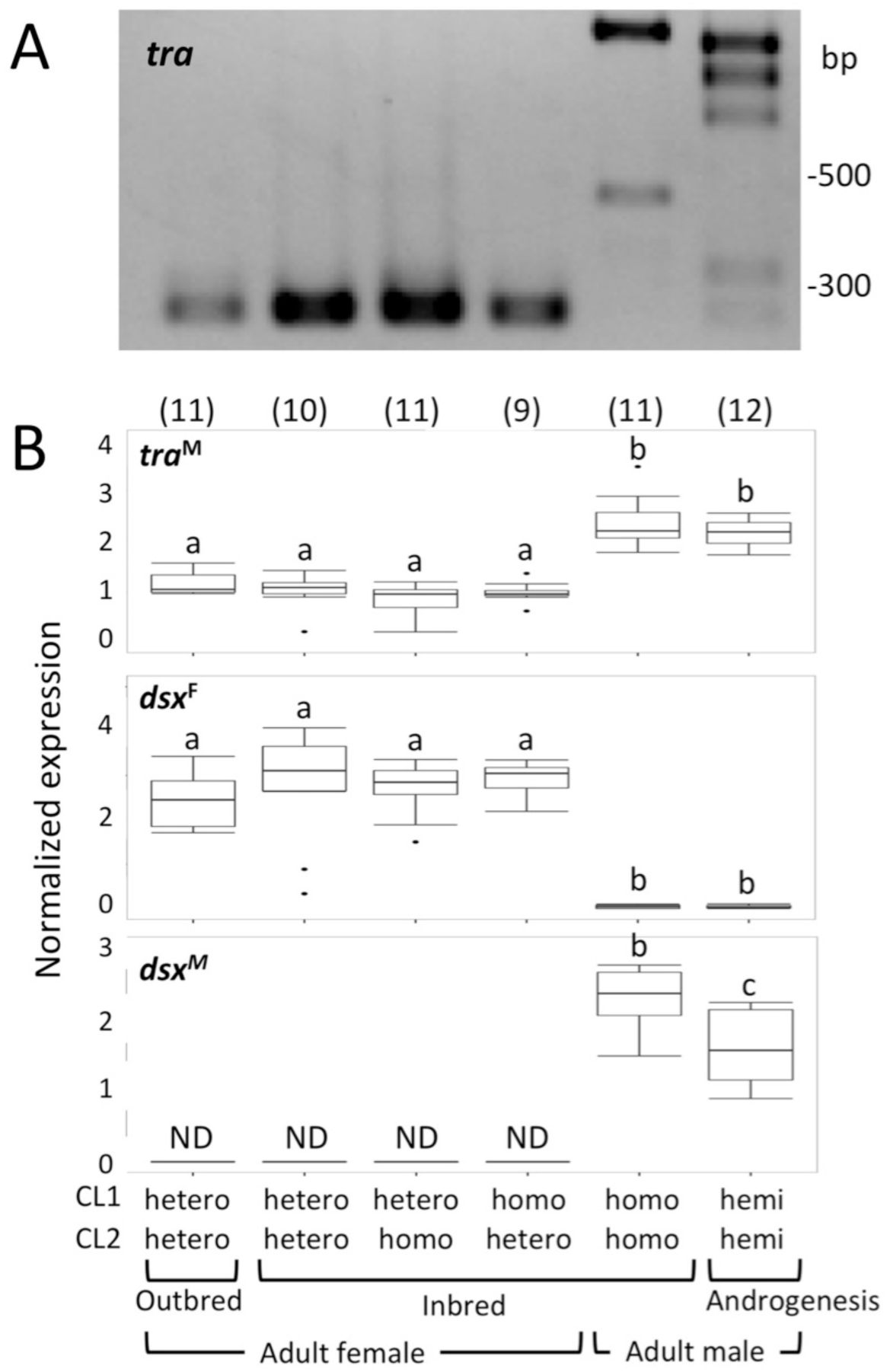
Sex-specific expression of *tra* and *dsx* correspond to the sexual phenotype. (A) Expression of female and male splice forms of *tra* in each allele type at two CSD loci. Splicing forms of other individuals are shown in *Supplementary figure 5- source data 4*. (B) Normalized gene expression (median ± SD) of *tra*^M^, *dsx*^F^ and *dsx*^M^ inferred by qPCR in each allele type at two CSD loci. Different letters indicate significant differences among groups (Tukey-Kramer test, P < 0.05). “ND” stands for “not detected”. Sample sizes are given in parentheses. Boxplots show the median and interquartile ranges (IQR) and 1.5 IQR. The following figure supplements are available for figure 6: **Supplementary figure 5.** Sex-specific splicing of *tra* corresponded to the sexual phenotype in the adult stage and allele types of two CSD loci *in V. emeryi*.

### Positive-regulatory-feedback loop at the level of *tra*

Gene expression data revealed that *tra*^F^ and *dsx*^F^ expression levels in females homozygous at one of the two CSD loci do not differ from those of females heterozygous at both CSD loci. As a mechanism for amplifying the initial signal derived from one heterozygous CSD locus, we hypothesize a positive-regulatory-feedback loop at the level of *tra*, as seen in *fem* of *A. mellifera* [10]. To test the existence of a positive-feedback loop of *tra* in *V. emeryi*, we injected *tra*^F^ mRNA with introduced synonymous substitution mutations (*tra*^F-mutORF^ mRNA) into embryos produced by inbreeding. After that, we assayed processing of endogenous *tra*^F^ mRNA in first instar larvae confirmed as males by HRM analysis (*Figure 7*). RT-PCR amplification showed that expression of *tra^F^* induces a switch from male to female mRNAs (*Figure 8A*-*source data 3*, *Supplementary Table 2*). Sequence analysis showed that *tra*^F^ mRNA detected in males was transcribed from the male genome, rather than derived from the injected *tra*^F-mutORF^ mRNA (*Supplementary figure 6*). On the other hand, males injected with GFP mRNA did not switch from male to female mRNAs (*Figure 8B*-*source data 3*). These findings suggest that injected *tra* activity trans-activates the endogenous *tra* gene. We conclude that expression of Tra protein induces its own synthesis by directing the processing of *tra* pre-mRNA to the productive female mode. This finding reveals that *tra*^F^ expression is amplified by a positive-feedback splicing loop in *V. emeryi*.

**Figure 7.**
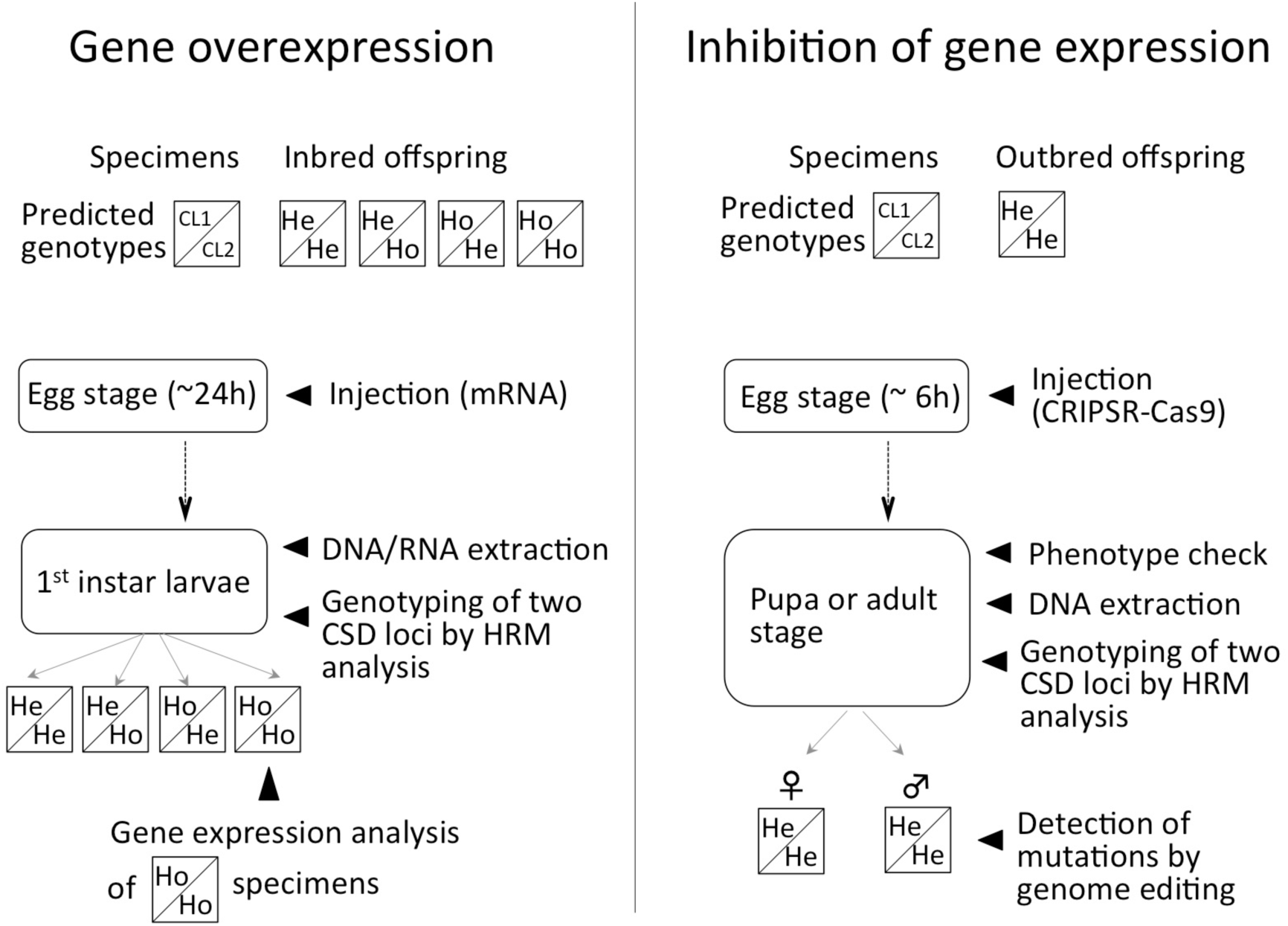
Experimental design for functional analyses described in the current study. For gene over-expression analysis, offspring produced by inbreeding crosses (25% of individuals should be homozygous at CL1 and CL2) were used. After mRNA injection, we genotyped all individuals grown to the 1st instar to select males that were homozygous at both CSD loci, and used their RNA for the latter gene expression analysis. To test the effect of inhibition of gene expression, we injected RNPs into outbred offspring produced during winter, since these are mainly female. After treatment, we checked sexual phenotypes of all individuals grown to pupae or adults, and confirmed their allele types at two CSD loci by HRM analysis. We conclude that phenotypic males, heterozygous at two CSD loci, result from sex reversal from female to male. In this figure, He and Ho indicate heterozygous and homozygous.

**Figure 8.**
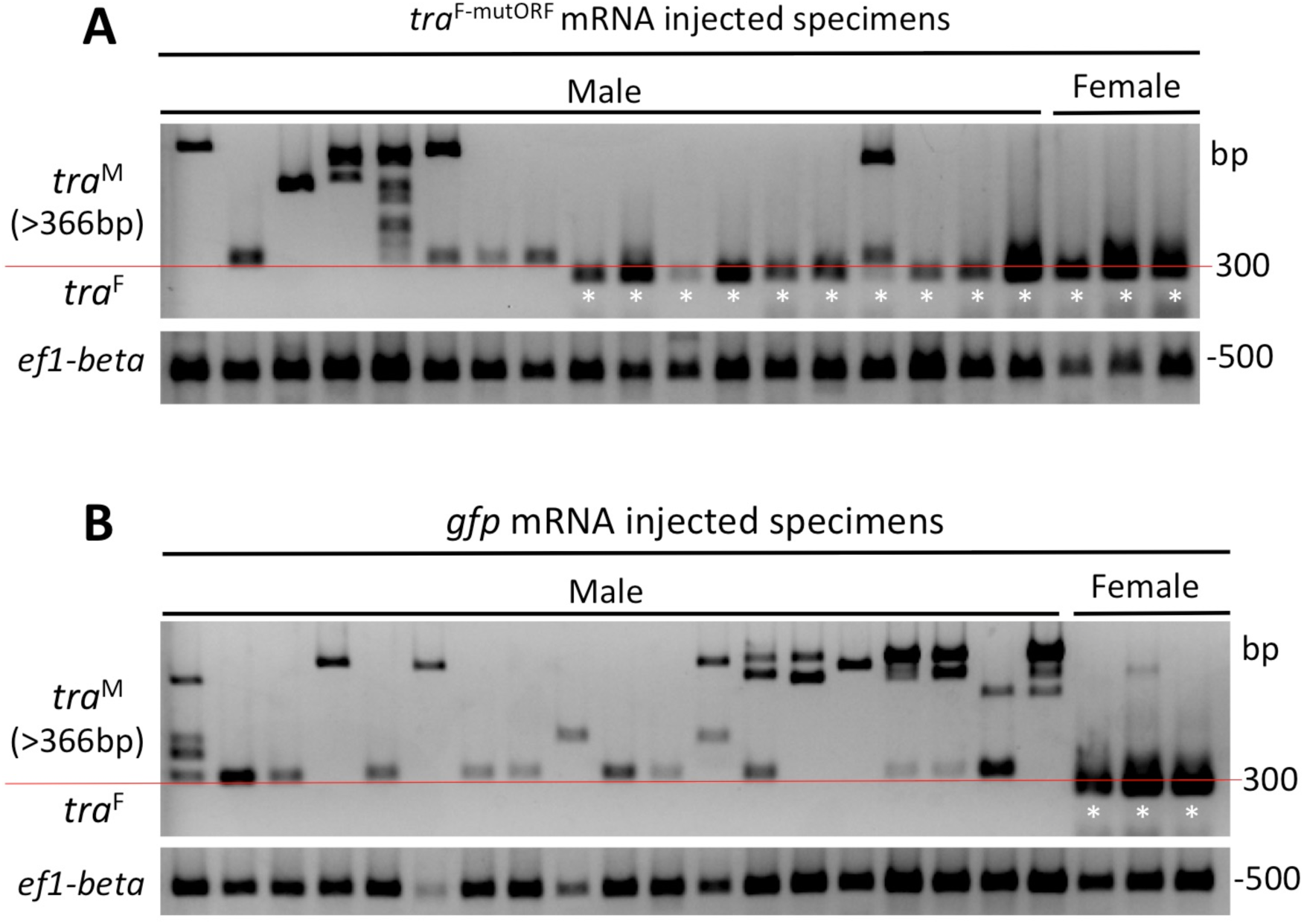
Processing of endogenous *tra*^F^ in response to injection of *tra*^F-mutORF^ mRNA in male embryos. (A) Ten of 18 males (first instar larvae) switched from male to female mRNA (277bp, fragments under 300-bp line with asterisk) by injection of *tra* introduced synonymous substitution mutations (*tra*^F-mutORF^) mRNA. (B) All 20 males injected with GFP-encoding mRNA showed only the male type of *tra* mRNA (≥ 326bp, fragments upper 300-bp line). In both conditions, “Females” comprised individuals of three allele types (CL1/CL2: hetero/hetero, hetero/homo, homo/hetero from left to right).

Using the same set of samples, expression of *dsx*^F^ and *tra*^M^ was tested by qPCR. Expression of *dsx*^F^ were detected in males with endogenous *tra^F^* expression after treatment of *tra^F-mutORF^ mRNA* although these are below the detection limit in males treated with *gfp mRNA* and males without endogenous *tra*^F^expression after treatment of *tra*^F-mutORF^ mRNA. However, expression level of *dsx*^F^ in males with endogenous *tra^F^* expression was significantly lower than in females injected with GFP (P < 0.05; *Supplementary figure 7*). Expression of *tra*^M^ in these males was significantly higher than that of females injected with GFP (P < 0.05; *Supplementary figure 7*), and did not differ from that of males injected with GFP. Although introduction of *tra*^F-mutORF^ mRNA in males induces splicing of endogenous *tra*^F^ (*Figure 8A*-*source data 3*), it does not affect expression of *tra*^M^, as described in honeybees [10]. During this experiment, no *tra*^F-mutORF^ mRNA-injected males ever developed into phenotypic female adults.

### Effect of *tra* inhibition

To investigate the effect of *tra* inhibition, we applied CRISPR/Cas9 to mutate *tra*^F^ and to express a nonfunctional Tra protein in early embryos (mainly female workers) produced by genetic-outbreeding, which are crosses using clonal queens (*Figure 2*a), and androgenetic males (*Figure 2*b) derived from the same colony. We performed zygosity tests on RNP-injected adults and pupae using HRM analysis (*Figure 7*). Although all individuals that showed heterozygosity at the two CSD loci were genetically female, 7 of 32 individuals showed phenotypic characteristics of wild-type males (*Figure 9*- *Supplementary figure 8*, *Supplementary Table 3*). For unknown reasons, all phenotypic males showed incomplete wing development and wing length varied among individuals. Sequence analysis showed deletion of the target genomic region with a frame shift in phenotypic males, but not in phenotypic females, suggesting that the male with a female genotype had changed sex from female to male due to the effect of *tra* inhibition. Our previous study revealed that sexual phenotypes correspond to the splicing patterns of *dsx* [37]. Thus, from this finding, we suggest that Tra protein directs splicing in *dsx* female mRNAs in *V. emeryi*.

**Figure 9.**
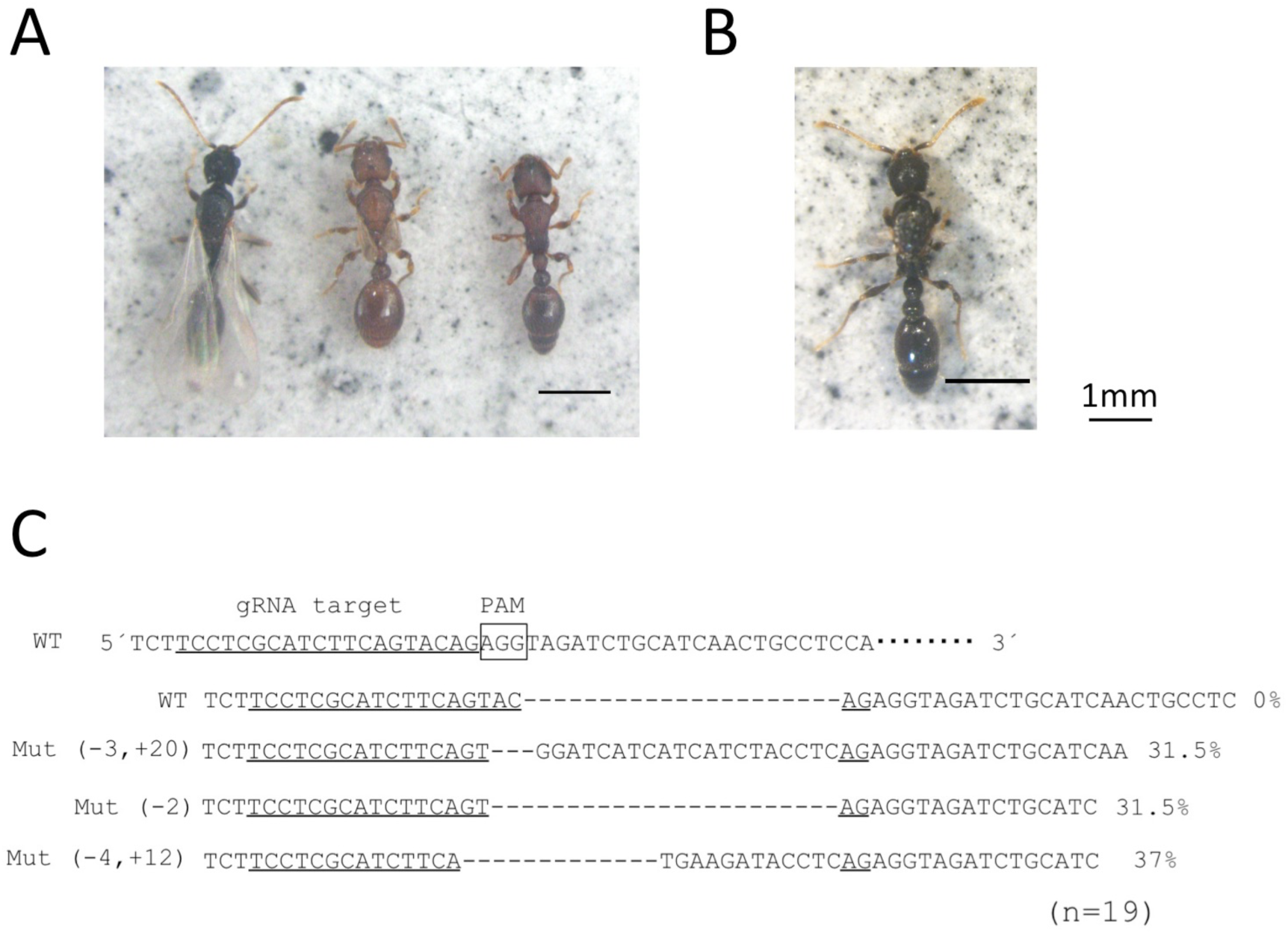
Detection of mutations in individuals that showed sex reversal from female to male after CRISPR-Cas9 mediation. (A) Wild-type phenotypes of male (left), queen (middle) and worker (right). (B) One of the seven phenotypic males that was heterozygous at two CSD loci. These individuals showed phenotypic characteristics of wild-type males, but with incomplete wing development. (C) Mutations detected in phenotypic males (B). The target site was in the third exon of the *tra* gene, which is located three nucleotides upstream of the PAM (AGG). The guide RNA (gRNA) target sequence is underlined. Nineteen PCR products for the third exon of *tra* show that this individual has at least three mutant alleles with frame shifts. Data of all individuals with sex reversal after treatment are shown in *Supplementary figure 8*. The following figure supplements are available for figure 9: **Supplementary figure 8.** Detection of mutations in individuals that showed sex reversal from female to male after CRISPR/Cas9 mediation.

### Allele types of CSD loci in parthenogenetically produced clonal queens

Gene expression and functional data using offspring produce by inbreeding showed that sex determination in *V. emeryi* relies on two CSD loci. On the other hand, the female determination system in clonal queens (*Figure 2*a,c) seems to be different [28]. In our previous study, we couldn’t find any evidences from genomic sequences of both *tra-like* at CL1 and the exonic regions at CL2 that clonal queens were hetelozygous at these loci, implying the existence of additional primary sex determination signals solely used for clonal queen production [28]. We applied HRM analysis to confirm zygosity state of two CSD loci in clonal queens and showed that they are homozygous C..T/C..T at CL1 and homozygous T/T at CL2. These facts support that a feminization of clonal queens is not controlled by two CSD loci, as predicted in previous study [28].

## Discussion

Our gene expression and functional analysis data revealed that functional Tra protein produced from *tra*^F^ is necessary for female differentiation in *ml*-CSD *V. emeryi*. We also suggest that a positive-feedback splicing loop of Tra could compensate for a shortage of Tra protein in individuals homozygous at one of two CSD loci, resulting in proper females. In *V. emeryi*, *tra*, located at the CL1 locus (*Figure 4*) is transcribed and expressed in sex-specific isoforms by alternative splicing. Both the sex-determination protein N-terminal and complementary sex-determiner protein regions were identified in female isoforms, but not in male isoforms in *tra* of *V. emeryi*, as with the *tra/fem* homologs of other holometabolous insects, including honeybees. Using samples from geographically separated sites, RT-PCR and expression (qPCR) analyses corroborated one another, strongly supporting the hypothesis that different isoforms and expression patterns are observed in the two sexes. Another *tra* homolog, *tra-like* (*Figure 4*), showed genomic structure similar to that of *csd* in honeybees. *tra-like* contains the sex-determination protein N-terminal domain, complementary sex-determiner protein and coiled-coil motif in both sexes, as with *csd* in honeybees [12,50,51]. In the last exon, numbers of repeated motifs differed among samples (*Supplementary figure 1AB*).

Although it is unclear whether the region actually reflects CSD allele specificity in *V. emeryi*, it might correspond to the hypervariable region (HVR) in *csd* of honeybees [50, 51]. An established zygosity test using HRM analysis enabled us to classify two CSD allele types of females produced by inbreeding and to compare allele types of two CSD loci and expression of genes related to the sex-determination pathway. Although females are homozygous at one of the two loci, *tra^F^* and *dsx^F^* expression did not differ from those of females heterozygous at both CSD loci.

The results of functional analyses suggest a mechanism to amplify the signal derived from heterozygous CSD, namely a positive-regulatory-feedback loop at the level of *tra*. We have demonstrated that introduction of female *tra* mRNAs induces female-specific *tra* mRNA splicing in males. We conclude from this observation that expression of *tra* establishes a positive-feedback loop in which Tra protein induces its own synthesis by splicing *tra* pre-mRNAs into the productive female mode, as concluded in *fem* of *A. mellifera* [10]. At the same time, this functional analysis also showed that introduced female *tra* RNA does not affect *tra*^M^ expression in male embryos, as shown in *fem* of honeybees [10]. In both *A. mellifera* and *V. emeryi*, introduced *fem^F^/tra*^F^ mRNA initiate positive-regulatory-feedback loop of *fem*/*tra*, but may not be sufficient to induce sex reversal from male to female, since primary masculinization signals are active in these males and splicing of *dsx* may not completely switch from male to female mode. On the other hand, sex reversal from female to male was induced by inhibition of *tra* in *V. emeryi*. We applied CRISPR/Cas9 to inhibit *tra* in early stages of female embryos, and induced phenotypic males, suggesting that *tra* mutations occurred almost at the single-cell stage and that Tra protein didn’t work during development. Given that sexual phenotypes correspond to splicing patterns of *dsx* in *V. emeryi* [37], our data indicate that Tra protein directs splicing in *dsx* female mRNAs.

Although a sex-determination system with multiple primary signals seems to be rare among diverse primary signals, it appears to have evolved founded on well-conserved base components of the sex-determination cascade and gene regulatory network [5,40,46]. Today, it is thought that positive feedback in the sex-determination cascade is the result of convergent evolution and is essential to arrive at a binary decision, as well as sex-specific splicing [46]. In holometabolous insects, genes involved in the sex-determination cascade, such as *tra/fem*, *Sex-lethal* (*Sxl*) in *Drosophila melanogaster,* and *IGF-II mRNA-binding protein* (*BmIMP*) in *Bombyx mori* are regulated by an autoregulatory positive-feedback loop [46,52,53]. In the parasitoid wasp, *Nasonia vitripennis*, with a CSD-independent sex determination system, the female-specific autoregulatory loop of *tra* is most likely regulated by maternal *tra* mRNA [21]. In addition, a CSD-independent factor, *wasp overruler of masculinization* (*wom*) is also needed to initiate female-specific *tra* expression [54]. Most conserved *tra*, and *sxl* in *D. melanogaster* directs the splicing *dsx* to female mode, while *BmIMP* in *B. mori* directs the splicing *dsx* to male mode [55]. Genes with a positive-feedback loop (*tra*, *fem* and *BmIMP*) located just upstream of *dsx* except *sxl* in *D. melanogaster* [56], convert an upstream signal into a binary response and direct splicing of *dsx*. In addition to providing a bistable system and maintaining the determined sex, *tra* in *V. emeryi* reduces multiple primary signals of the same hierarchy to two, enabling proper sexual differentiation. On the other hand, in *L. fabarum*, with four-unlinked CSD loci, as in *Athalia rosae*, there was no *tra* homolog in any of these regions and it is unlikely to be in other genomic regions [30, 57]. Given that the CSD-based sex-determination system works, *L. fabarum* may have a novel intermediate gene instead of *tra* that amplifies the signal derived from heterozygous CSD and exceeds the required expression threshold for feminization.

Our previous expression analysis suggested that two primary sex determination signals are integrated into *dsx* [37]. Using inbred offspring with each allele type of two CSD loci, our current study further suggests the following: 1) The two primary sex determination signals are integrated into the upstream of *dsx*, namely *tra*. 2) The primary signal derived from heterozygous CSD is amplified by an autoregulatory feedback loop of *tra*. 3) *tra* directs splicing of *dsx* (*Figure 10*). Based on the present results, we hypothesize a sex determination cascade with a positive-feedback loop of *tra* (*Figure 10*). In adults, expression of *tra*^M^ and *dsx*^M^ mRNA in females homozygous at one of the two CSD loci was not significantly different from that of females heterozygous at both CSD loci (*Figure 6*), suggesting that most pre-mRNAs are allocated to female isoform synthesis (*Figure 10*). This model system, amplifying heterozygous CSD signals, allows maintenance of a female hetero, male hemi/homo sex-determination system regulated by multiple primary signals. In order to fully understand the sex-determination cascade in *V. emeryi*, it is necessary to identify the two primary signals that directly or indirectly splice *tra*. So far, *tra-like* is the most likely candidate gene, working as one of the two primary signals, since it is located near the QTL-related markers in CL1 (*Figure 4*) [28] and shows domains and motif similar to *csd* of *Apis* species (*Figure 3A*-*Supplementary figure 1*). In contrast to the idea that *ml*-CSD emerged from duplications of an ancestral single-locus system [26, 27], primary signals located in CL2 may not be homologs of *tra* [28]. In each candidate encoded in CL1 and CL2, identification of the region reflected by the CSD allele specificity, e.g., HVR in honeybees [50, 51], is required to confirm that they act as primary sex determination signals.

**Figure 10.**
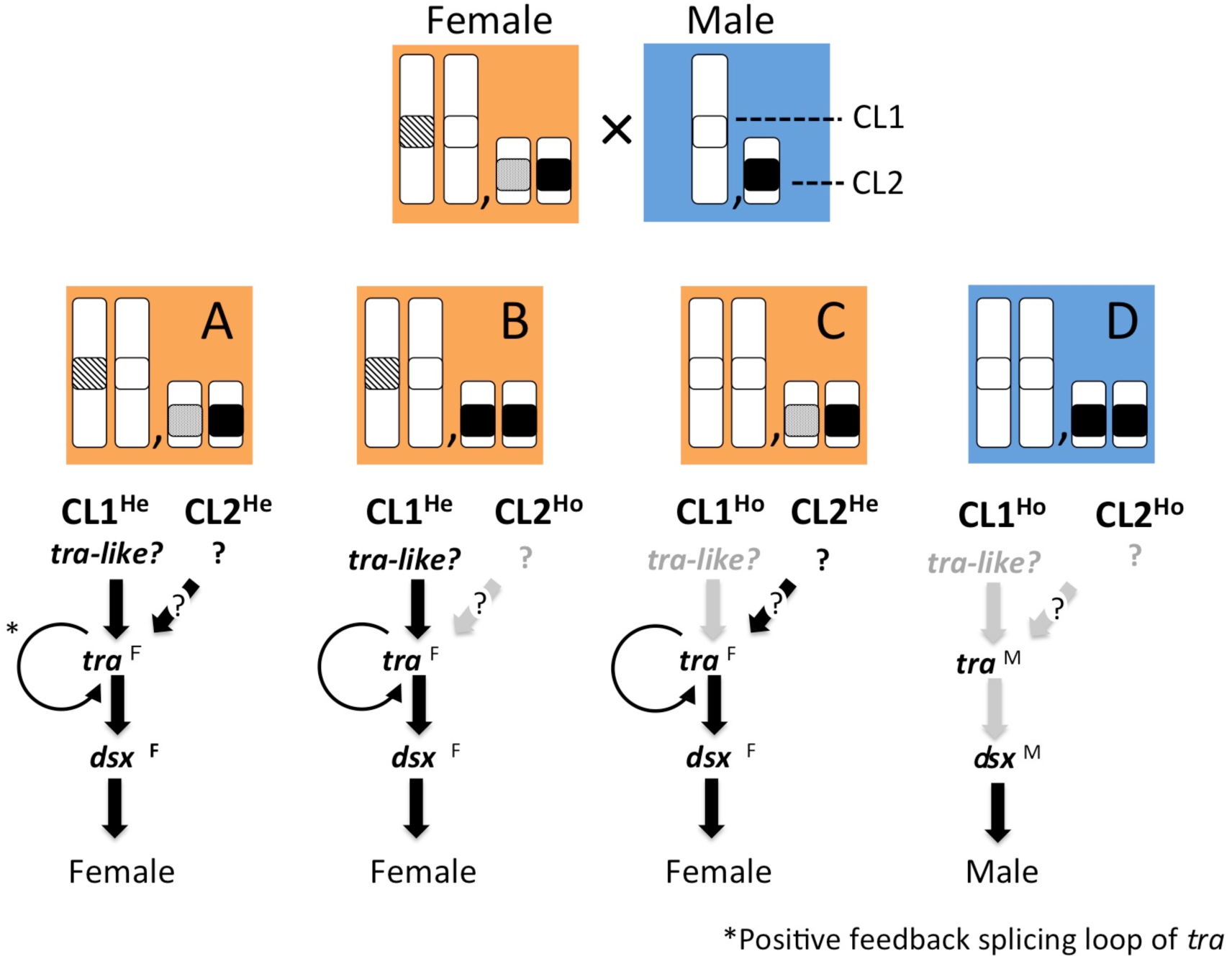
Predicted sex determination cascades with a positive-feedback-splicing loop of *tra*. We expected a signal pathway in which the primary signal from CL1 (so far, the most putative gene is *tra-like*) regulates *tra* and *dsx*. Another primary signal from CL2 is also integrated into *tra*. The signal derived from the heterozygous allele is amplified by the positive-feedback-splicing loop of *tra.* As with females with heteroallelic CL1 and CL2 (A), individuals with heteroallelic CL1 and homoallelic CL2 (B) or vice versa (C) show female development since signals derived from heteroallelic alleles are sufficiently amplified by *tra* to splice *dsx* into female mode. (D) Individuals with homoallelic CL1 and CL2 show male development. In this case, *dsx* is spliced into the male mode since no signals are produced from either CL1 and CL2.

The sex determination system in sexually produced females in *V. emeryi* relies on two CSD loci; however, the feminization mechanism of clonal queens could not be explained by zygosity state of two CSDs. HRM analysis showed homozygous at both CSD loci, as with previously predicted by genomic sequencing data of *tra-like* and exonic region at CL1 and CL2, respectively [28] (*Figure 2*). These fact suggest that feminization of clonal queens might be initiated by a CSD-independent factor (e.g., maternal provision of *tra*^F^ mRNA [21, 58], CSD-independent zygotic factor which initiate *tra*^F^ expression [54], epigenetic factor directing female development). Further studies will be needed to resolve the whole feminization system in *V. emeryi*, and the mechanism to sustain two separate sex determination systems in one species. The existence of two sex determination systems in one species, depending on life-history stages, may provide interesting insights into evolution of sex determination systems.

Today, molecular mechanisms of sex determination in the Hymenoptera have not been clarified, except in a few model species such as *A. mellifera* and *N. vitripennis* [10–12,21]. Comparative genomic studies have noted that tandem copies of *tra*, as with *csd*/*fem* in honeybees, occur in many ants [28,38,39,59]. However, without functional studies, these data have limitations in reference to the evolutionary origin of these sex determination genes. It is still unclear whether the duplication of *tra* has an independent evolutionary origin within the Aculeata, or whether it occurred before the divergence of this group. Therefore, so far, we cannot conclude that *tra-like* and *tra* in CL1 in *V. emeryi* (*Figure 4*) correspond to *csd* and *fem* in honeybees, respectively. To address the issue of evolutionary trajectories of these genes, functional data regarding sex-determining genes in a wide variety of ants is essential. Now *V. emeryi* is becoming a powerful model ant species with methods for functional analysis, such as genome editing, in addition to ease of rearing and inducing breeding in the laboratory. These established methods should be of great help in advancing studies to identify the molecular sex determination system of *V. emeryi* in the future.

## Materials and Methods

### Sample preparation

*V. emeryi* colonies were collected from Tokyo, Tochigi, and Ishikawa Prefectures in Japan from April to October 2016 and have been kept in the lab. These collected colonies were maintained in artificial plaster nests at 25 °C under a 16/8 h light/dark cycle. Colonies were provided with tap water, dry crickets, EBIOS tablets (Asahi Corporation Co., Ltd.) and brown sugar every other day. We sampled emerging workers and new reproductive individuals in field and laboratory colonies for later experimental crosses and molecular experiments.

### Experimental crosses

We conducted experimental crosses to produce new offspring for laboratory studies. *Vollenhovia emery* has evolved an extraordinary reproductive system that involves both parthenogenesis and androgenesis (*Figure 2*) [33, 34]. Briefly, almost all queens are clones of their mothers (parthenogenetic clonal queens), and haploid males are clones of their fathers (androgenetic clonal males) (*Figure 2*). Sterile workers and a few queens (sexual queens, *Figure 2*d) develop from fertilized diploid eggs [36]. In this study, crosses using sexual queens and androgenetic males are denominated “inbreeding”, which is equivalent to a backcross (*Figure 2*). In contrast, crosses between clonal queens and androgenetic males, which naturally occur in the field, are termed “genetic-outbreeding”, since they never share genomes (*Figure 2*). All of these samples produced by laboratory crosses were subjected to RT-PCT, qPCR, and functional analysis.

### Cloning full-length *tra* and *tra-like* genes

Previous QTL analysis showed that one of the two CSD loci (CL1) contained two tandem copies of *tra* homologs (*traA* and *traB, NCBI NM_001310035.1 and NM_001310036.1*) on the same scaffold [28] (*Figure 4*). According to their genomic organization and gene expression manner, we refer to *traA* and *traB* as *tra-like* and *tra* (*Figure 4*). Both *tra* homologs contain the sex-determination protein N-terminal (pfam12278) and complementary sex-determiner protein regions (pfam11671), as with honeybees and other ants. We therefore determined the full-length sequences of *tra-like* and *tra* using RACE to identify these isoforms (NCBI accession LC677157-LC677165). Total RNA was extracted from 15 queens, 15 workers, and 15 males using ISOGEN II reagent (Nippon Gene Co., Ltd.). These female specimens were collected from a site in Ishikawa Prefecture. In our previous study, QTL analysis showed that these females have two sex-determination loci [28]. After DNase treatment using a TURBO DNA-free Kit (Ambion), 1.5 µg of total RNA of each sex were subjected to 5’ and 3’ RACE with a GeneRacer kit (Invitrogen) according to the manufacturer’s instructions. PCR primers for the RACE assay are given in *Supplementary Table 1*. Amplified products were purified using NucleoSpin Gel and PCR Clean-up (Takara Bio), and cloned into the pGEM-T easy vector (Promega). After determining the nucleotide sequences using a 3500 Genetic Analyzer (Applied Biosystems), they were aligned to *V. emeryi* genomic DNA using Splign [60] to identify exon-intron boundaries. Based on the Splign results, we then designed specific PCR primers to detect each exon and sex-specific splicing variant (*Supplementary Table 1*).

### Real-time quantitative PCR

For the qPCR assay to compare *tra* expression between sexes, we used samples collected from all three sites. Total RNAs of males and females were extracted from 4 to 7 pupae and 4-8 adults using ISOGEN II reagent (Nippon Gene Co., Ltd.), with three replicates prepared for each sample. Then, 1 µg of each total RNA was subjected to RT-PCR using a High-Capacity cDNA Reverse Transcription Kit (Applied Biosystems).

To compare *tra* and *dsx* expression among individuals with each allele type at two CSD loci, 30 females and 11 diploid males produced by inbreeding, and 12 androgenetic haploid males were used. Total RNAs and DNAs were extracted using ISOGEN II and ISOGENOME reagents (Nippon Gene Co., Ltd.), respectively. In this analysis, specimens derived from Tokyo Prefecture were used. After DNase treatment using a TURBO DNA-free Kit (Ambion), RNAs that had been purified and concentrated using NucleoSpin RNA Clean-up XS (Takara Bio) were reverse-transcribed using a High-Capacity cDNA Reverse Transcription Kit (Applied Biosystems).

Real-time PCR and statistical analyses of *tra*^M^, *dsx*^F^ and *dsx*^M^ expression were conducted using the same method as in our previous study [37]. Since primers for qPCR could not be designed for the *tra*^F^ sequence, RT-PCR comparison was performed between the male and female isoforms with a primer pair designed to amplify the sex-specific exon of *tra*. A housekeeping gene, *ef1-beta*, was used as an internal control for RNA extraction and qPCR [37]. Primers for RT-PCR and qPCR are listed in *Supplementary Table 1*.

### High-resolution melting curve (HRM) analysis

Female hetero- and male homozygous genomic mutations in the two CSD QTL regions (CL1 and CL2) were visually searched using the Integrative Genomics Viewer [61] with *V. emeryi* genomic and Rad-seq data [28]. We designed primers in *NCBI NW_011967057.1* and *NCBI NW_011967112.1*, which should link to the sex-determination genes in CL1 and CL2 (*Figure 4*). To evaluate whether the HRM assay is applicable to zygosity testing of the two CSD loci, we confirmed that females produced by inbreeding were heterozygous for least one of the two CSD screening regions, and that diploid males were homozygous at both regions. In this experiment, DNA was extracted from whole adults using 5% Chelex 100, as described previously [35]. After that, DNAs (1 µL of each) were used directly in 11.7 µL reaction volumes containing 1 µL of EvaGreenDye (Cosmo Bio Co., Ltd.), 2.5 µL of 5×PrimeSTAR GXL Buffer, 0.25 µL of PrimeSTAR GXL DNA Polymerase, 1 µL of 2.5 mM dNTP, 0.5 µL of each primer (10 µM forward and reverse primers), and 4.95 µL of RNAse-free water. PCR amplification, DNA melting, and end-point fluorescence level acquiring PCR amplifications were performed on a LightCycler 480 system (Roche Life Science). PCR amplification conditions were as follows: 95°C (10 min), then 48 cycles at 95°C (10 s) / 60°C (10 s) / 72°C (10 s). High-resolution melting analysis was performed after PCR amplification, that is, temperature ramping from 65 to 95^◦^ C, rising by 0.02 ^◦^C/s. Melting curves were normalized between two normalization regions before and after the major fluorescence decrease, representing melting of the PCR product, using LightCycler 480 Gene Scanning Software. Zygosity detected by HRM analysis was confirmed by direct sequencing using the same DNA samples chosen randomly. Primers for HRM analysis and direct sequences are listed in *Supplementary Table 1*.

Finally, to assess the accuracy of regulation of the two-locus sex-determination system at genetically independent sites, we conducted HRM analysis using females produced by inbreeding. Total numbers of tested females were 139, 123, and 126 from Ishikawa, Tochigi, and Tokyo, respectively. In addition, allele types of two CSD loci using 25 diploid males at each site were determined. After DNA extraction, HRM analysis was performed as described above. To test whether the proportions of females with each allele type are equal, Pearson’s chi-squared test was performed using R (version 3.1.1) (R core team).

### Flow cytometry for DNA quantification

To confirm the relationship between ploidy level and allele patterns of two CSD loci, we conducted flow cytometry using adult males and females produced by inbreeding crosses. Samples were prepared using a CyStain UV Precise P Kit (Sysmex Partec., GmbH.). Each body, except the abdomen was chopped with microscissors in nucleus extraction buffer, stained for 15 min in a staining buffer, and filtered with a CellTrics Filter using a mesh diameter of 30 µm (Sysmex Partec., GmbH). Androgenetic males (which are expected to be haploid) were also used for the analysis to compare DNA quantities between haploid and diploid individuals. DNA contents of dissociated cells were measured with a Cell Analyzer EC800 (Sony Corp). At the same time, each abdomen was used for HRM analysis to test the allele types of two CSD loci. We prepared at least two samples for each allele type of two CSD loci in addition to two androgenetic haploid males for this experiment.

### Gene Overexpression

To introduce *tra*^F^ mRNA synonymous-substitution mutations (*tra*^F-mutORF^ mRNA), twelve nucleotides at the *tra*^F^ ORF were substituted. Replacement of nucleotides was conducted with a PrimeSTAR Mutagenesis Basal Kit (Takara Bio) using full length of *tra*^F^ mRNA encoding plasmid DNA as a template (see primers at *Supplementary Table 1*). Linear templates with the T7 polymerase promoter sequence at the 5′ end were prepared by PCR using primers shown in *Supplementary Table 1*. The mRNA (*tra*^F-mutORF^ mRNA) was generated using the mMESSAGE mMACHIN T7 ULTRA Transcription Kit (Thermo Fisher Scientific) in which the 5′ cap structure and Poly (A) tail were added during RNA synthesis.

For microinjection preparation, 0–24-h embryos produced by inbreeding were transferred and aligned on a double slide attached with tape to a microscope slide using fine forceps. Our cytological observations suggest that embryos of ∼24h are the syncytial stage in which nuclei divide without cytokinesis. Next, eggs on a microscope slide were desiccated by placing them in a sealed box with silica gel (Nacalai Tesque., Inc.) for 30 min. Synthesized mRNA (0.08 pg) was injected from the long axis side of the embryo under a stereo microscope using a FemtoJet 4i injector (Parker Hannifin Corp). Then, embryos were transferred into a plaster nest with 20 workers for 10 days until the embryos had just hatched into first instar larvae. RNA and DNA were extracted from the larvae using a NucleoSpin RNA XS and a NucleoSpin RNA/DNA Buffer Set (Takara Bio). If individuals were homozygous at two CSD loci (male genotype) as a result of HRM analysis, their RNA was reverse-transcribed. Endogenous *tra*^F^ mRNA synthesis was checked with primers (*Supplementary Table 1*) that amplify the sex-specific exon of *tra* (female 277 bp, male ≥ 326 bp), but almost not *tra*^F^ mRNA-introduced synonymous substitution mutations (*tra*^F-mutORF^ mRNA) (*Supplementary figure 9-source data 5*). We also checked the sequence of cDNA of *tra*^F^ amplified in male injected *tra*^F-mutORF^ mRNA to investigate whether *tra*^F^ is transcribed from the male genome, not derived from injected mRNA. Functional analysis was performed using samples from Tochigi and Tokyo Prefectures.

### CRISPR/Cas9

CRISPR/Cas9 single-guide RNAs (sgRNA) that target Tra were designed with CRISPRdirect software [62]. sgRNA DNA templates were transcribed *in vitro* using a GeneArt Precision gRNA Synthesis Kit (Invitrogen) according to the manufacturer’s protocol. Primer and guide RNA sequences are in *Supplementary Table 1*. The resultant sgRNAs and Cas9 protein (TrueCut Cas9 Protein v2, Invitrogen) were diluted in RNase-free water to final concentrations of 500 and 600 ng/µL, respectively. These ribonucleoprotein RNPs were stored at −80°C until use. For microinjection preparation, 0–12-h embryos produced by genetic-outbreeding (*Figure 2*), which were mainly worker-destined embryos, were used. Microinjection was performed as described above. Injected embryos were maintained with about 50 workers until the embryos had developed into pupae or adult, which is the developmental stage at which sexual phenotypes are easily identified. After phenotypic observation, genotypes of both CSD loci were determined by HRM analysis. The target genomic region was amplified with PrimeSTAR GXL DNA Polymerase (Takara Bio) using a set of primers (*Supplementary Table 1*). After adding a 3′ dA overhang, resulting fragments were subcloned into the pGEM-T Easy vector (Promega) according to the manufacturer’s instructions. At least 8 clones for each sample were sequenced using the Sanger method that included Big Dye terminator Ver. 3.1 (Thermo Fisher Scientific) on an ABI 3730 Genetic Analyzer DNA sequencer (Applied Biosystems). We assumed that individuals with male phenotypes and female genotypes (CL1: hetero / CL2: hetero) had changed sex from female to male by genome editing.

## Acknowledgments

We thank members of the Laboratory of Environmental Physiology for discussions. We are grateful to Steven D. Aird for editing and comments on the manuscript. This study was supported by a Japan Society for the Promotion of Science (JSPS) Research Fellowship for young scientists (201940074).

## Author contributions

Misato Okamoto Miyakawa, Conceptualization, Methodology, Funding acquisition, Investigation, Visualization, Writing-original draft, Project administration, Writing-review and editing; Hitoshi Miyokawa, Methodology, Project administration, Writing-review & editing.

## Supplementary Information

**Supplementary figure 1.**
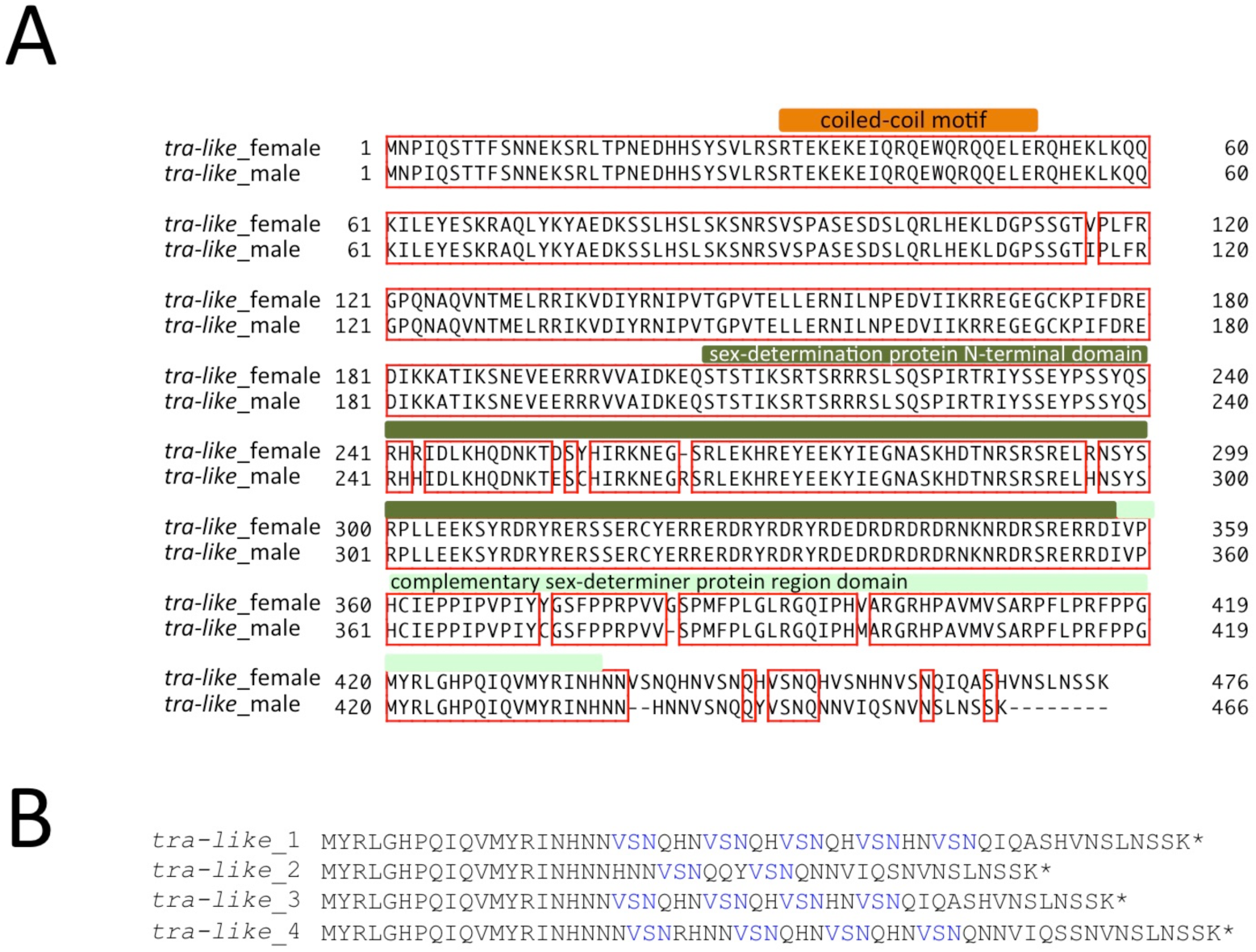
cDNA and amino acid sequence of *tra-like* in *V. emeryi*. (A) The amino acid sequence of *tra-like* isolated from females showed five repeats of VSN, whereas that of males showed two repeats at the end of the complementary sex-determiner protein region domain. Predicted domains and motifs are highlighted. (B) Alignment of the amino acid sequence region containing a repeated motif (VSN, HQN, etc.). So far, four types of sequences with different numbers of amino acid repeats have been isolated (*tra-like_*1: clonal queens in Ishikawa, Tochigi and Tokyo, *tra-like_*2: androgenetic males in Ishikawa, *tra-like_*3: androgenetic males in Tochigi, *tra-like_*4: androgenetic males in Tokyo).

**Supplementary figure 2.**
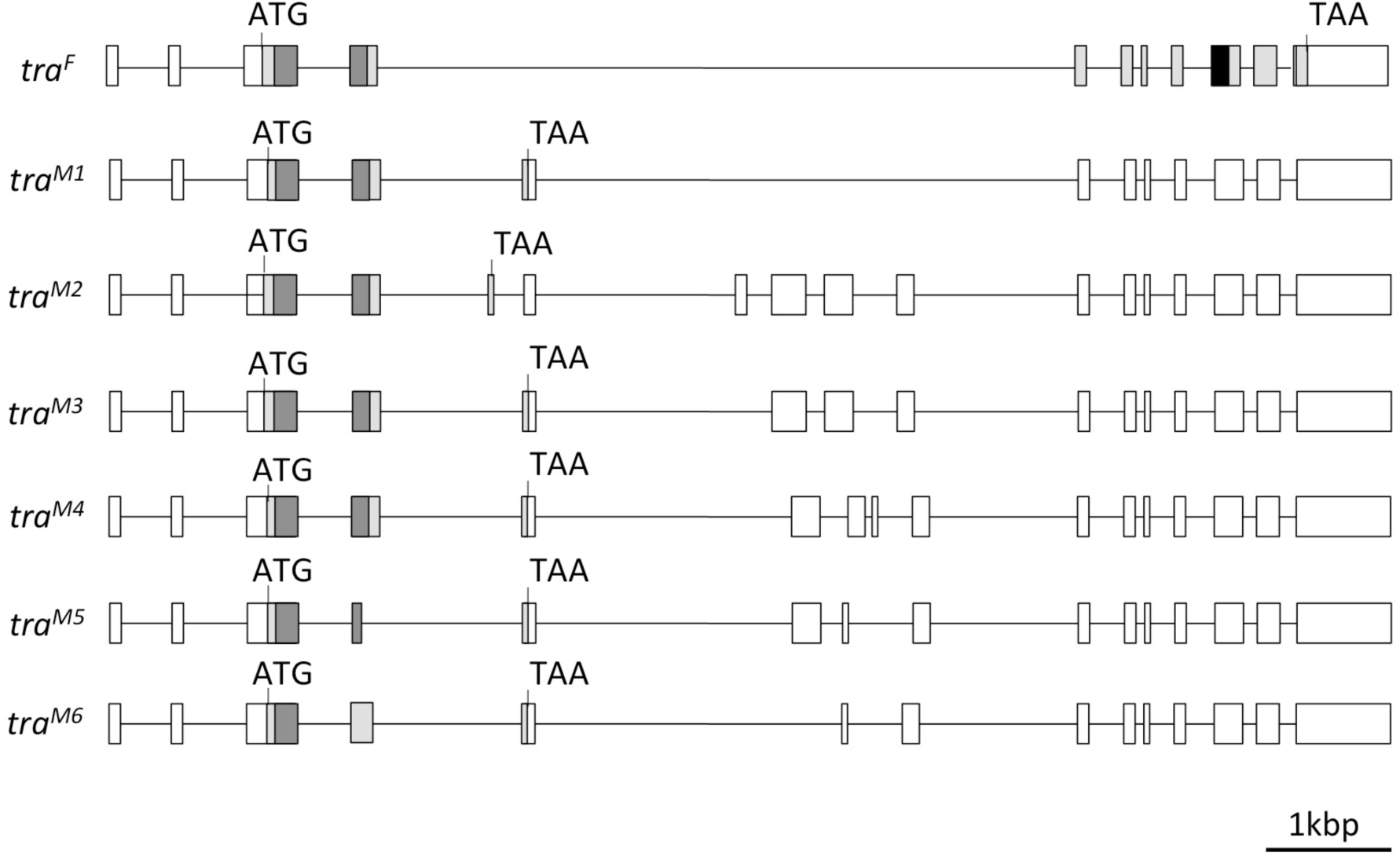
Exon (boxes) and intron (lines) structure diagram of *tra* in *V. emeryi*. In contrast to the female isoform (*tra*^F^), all six detected *tra* male isoforms (*tra*^M^) show an early termination codon at exon 4 or 5, and male-specific exons occur within the region corresponding to the intron in *tra*^F^.

**Supplementary figure 3.**
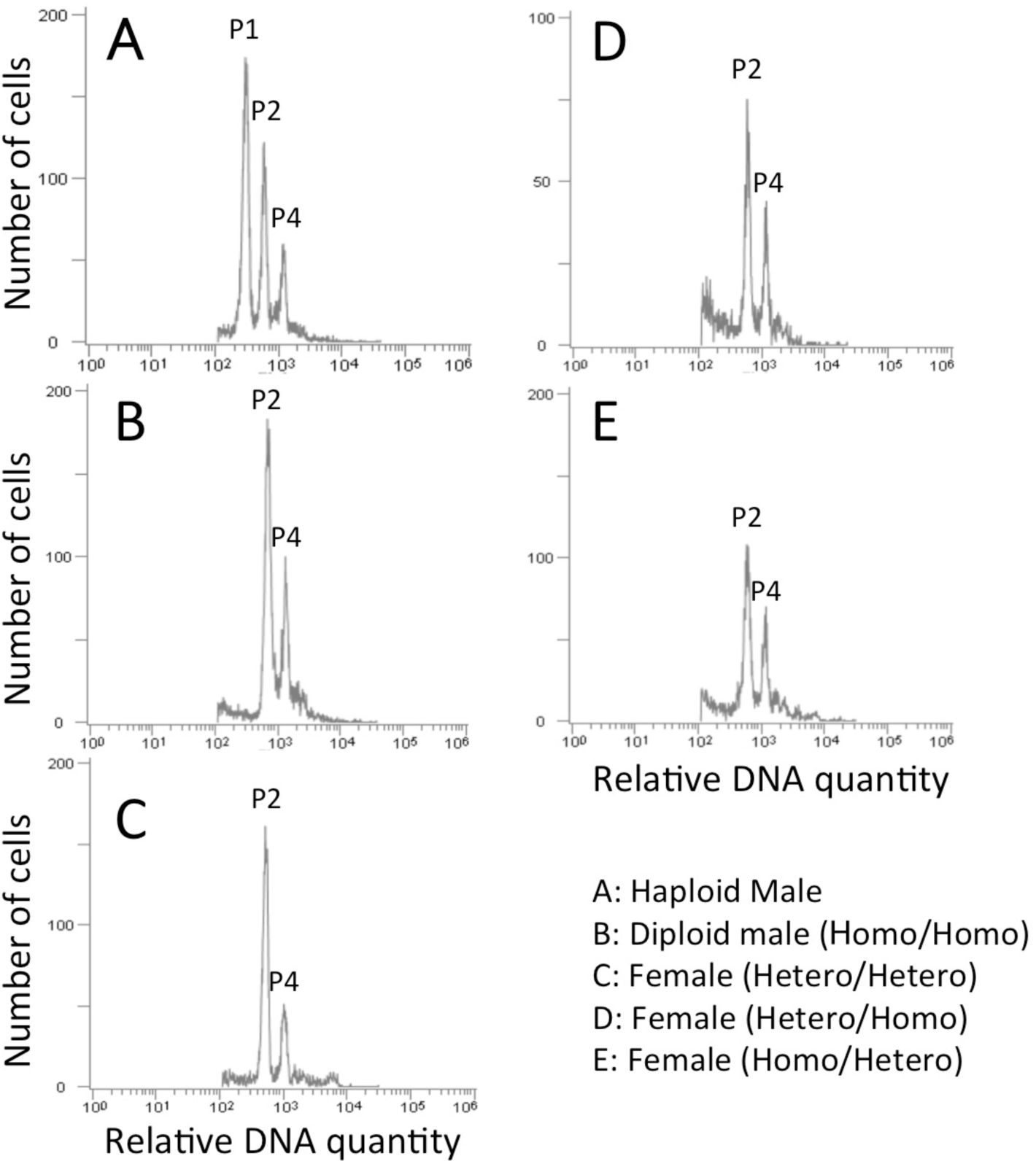
Flow cytometric DNA histogram in an androgenetic haploid male (A) and offspring produced by inbreeding crosses (B-E) in *V. emeryi*. Each histogram shows the cell frequency with regard to DNA content for a single individual. Haploid males and inbred offspring with each allele type of two CSD loci showed peaks in cell numbers according to their expected relative differences in DNA quantity. This result suggests that all inbred offspring, including males, are diploid. In haploid males (A), P1 and P2 peaks correspond to haploid nuclei and twofold haploid nuclei before cell division. The P4 peak corresponds to the distribution of polyploid cells. In diploid males (B) and females (C-E), Peaks P2 and P4 correspond to diploid nuclei and twofold diploid nuclei before cell division. We prepared at least two samples for each condition, and showed one of these results.

**Supplementary figure 4.**
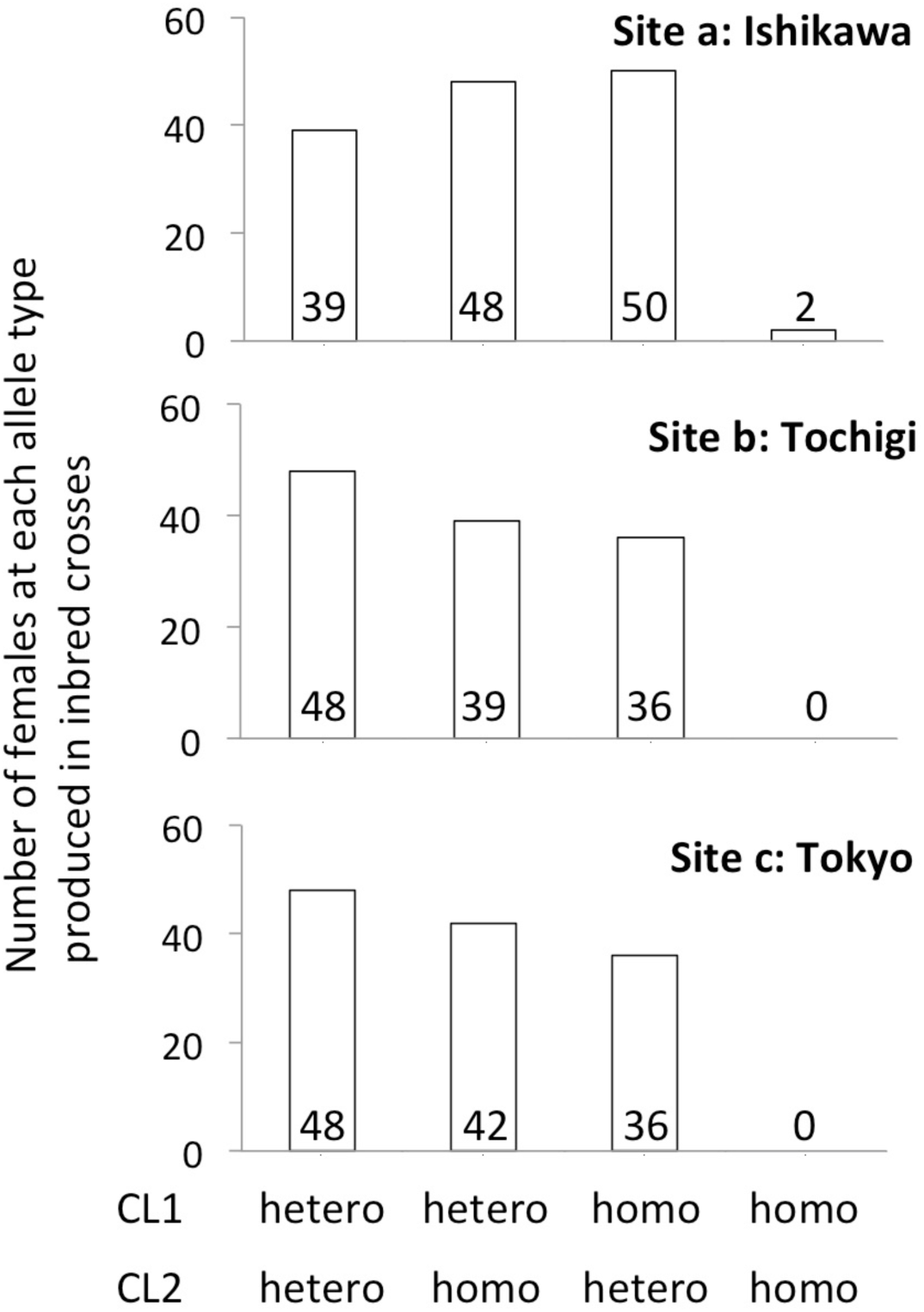
Almost all females produced in inbred crosses were heterozygous for at least one of the two CSD loci. Proportions of each three-allele type (CL1/CL2: hetero/hetero, hetero/homo, homo/hetero) in these females were statistically equal in all experimental sites (Pearson’s chi-square). These data suggest that factors other than two-loci CSD system, e.g., maternal factor, are not involved in sex determination in these females. Numbers of individuals tested by HRM analysis are shown in each box.

**Supplementary figure 5.**
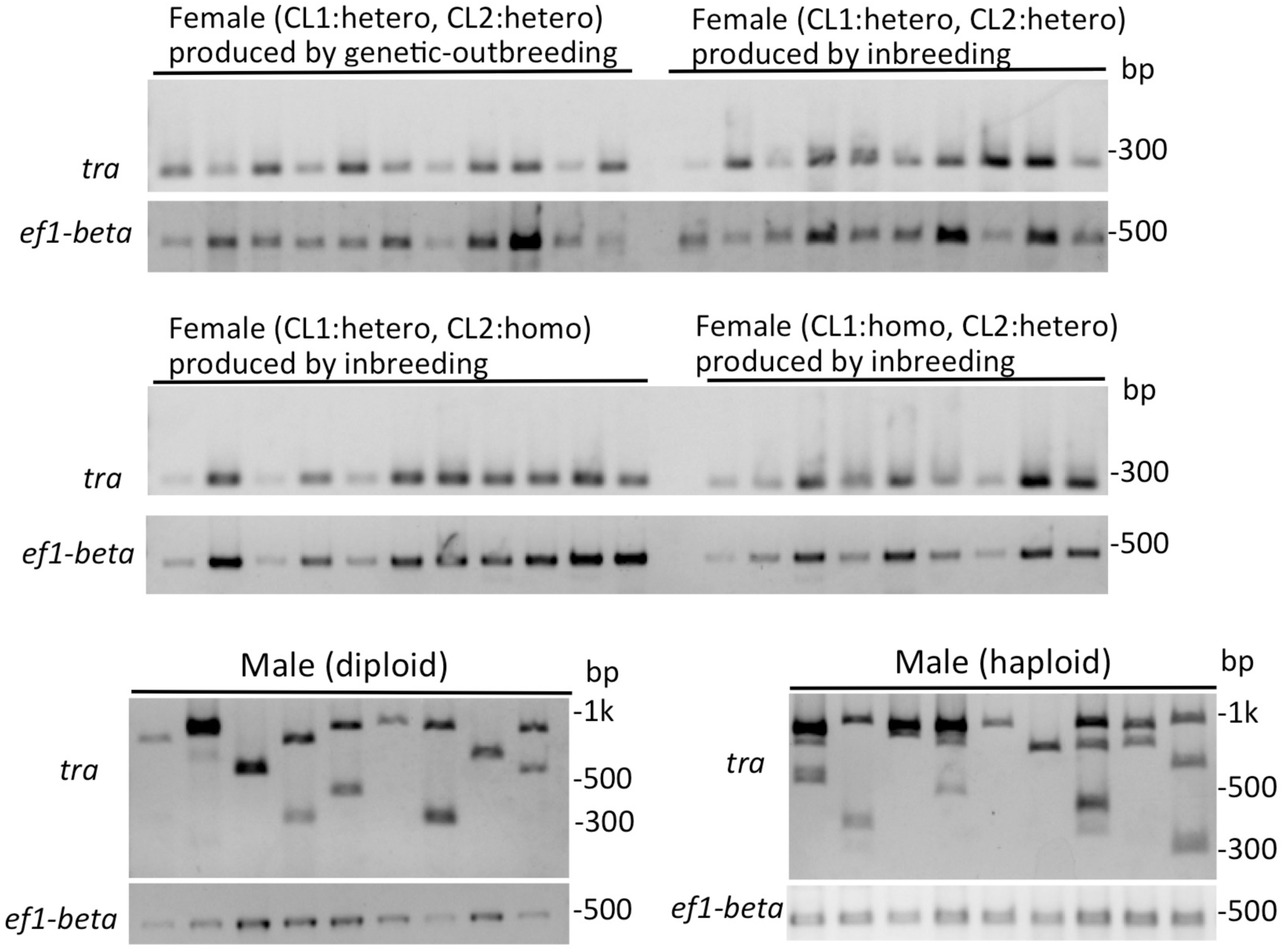
Sex-specific splicing of *tra* corresponded to the sexual phenotype in the adult stage and allele types of two CSD loci *in V. emeryi.* Irrespective of alleles at two CSD loci (CL1/CL2: hetero/hetero, hetero/homo, homo/hetero) and crossing type (genetic-outbreeding or inbreeding, Figure 2), all adult females show female *tra* transcripts (277 bp). On the other hand, all males show male *tra* transcripts (≥ 326 bp) at two CSD loci, irrespective of ploidy level (haploid hemi or diploid homozygous). All samples were used for the expression analysis shown in Figure 6, but two diploid and three haploid male specimens were not successfully amplified by PCR.

**Supplementary figure 6.**
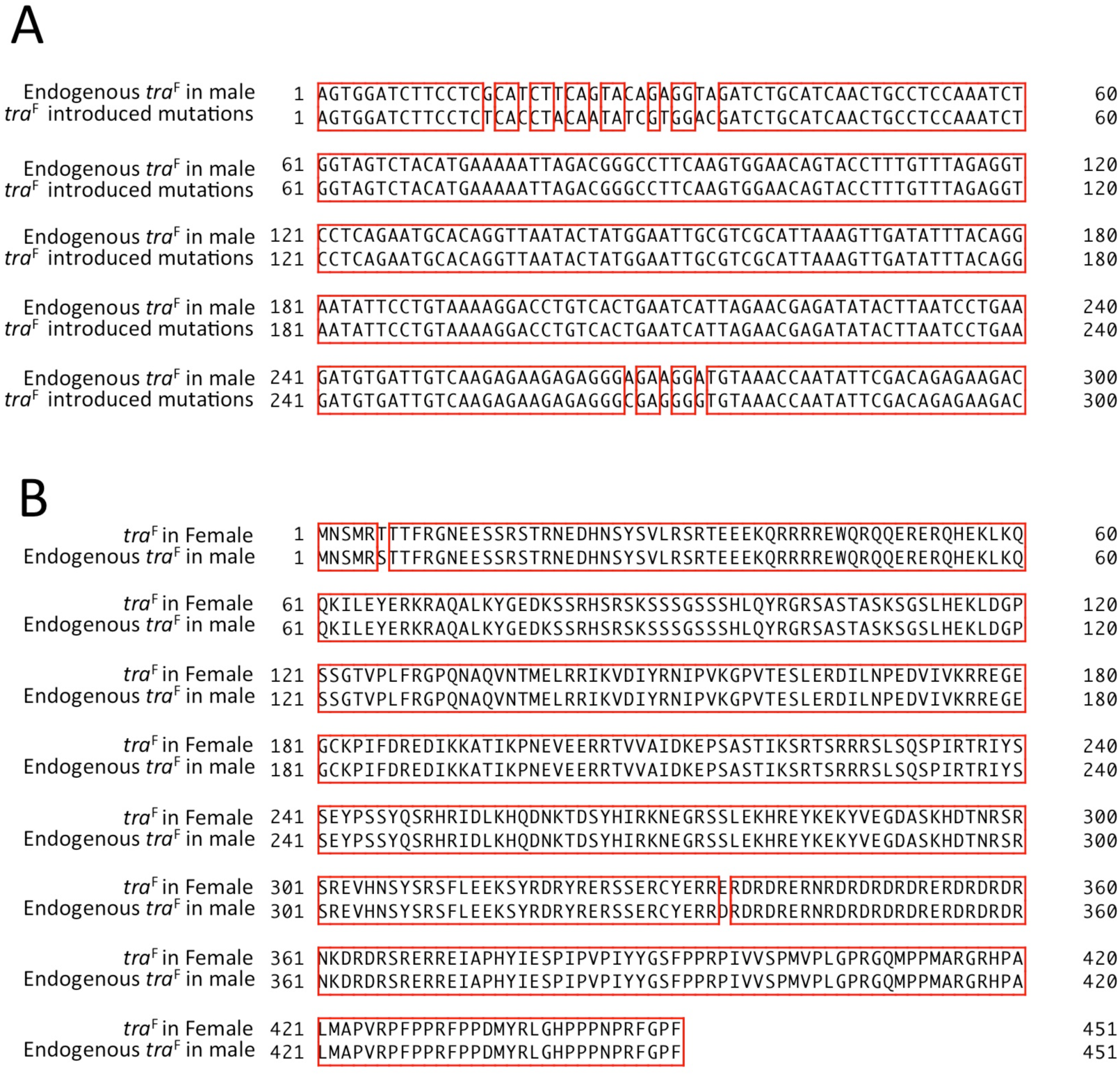
cDNA and amino acid sequences of *tra*^F^ synthesized in males by gene overexpression. (A) Comparison of cDNA sequences between injected *tra*^F^ and endogenous *tra*^F^. Introduced mutations were not detected in *tra*^F^ synthesized in males by gene overexpression. (B) Comparison of amino acid sequences of the coding region in *tra*^F^ synthesized in males and in wild-type females. Although two non-synonymous substitution mutations were detected between the sexes, amino acid properties of these mutations were conserved: T (threonine) and S (serine) are polar amino acids while E (glutamic acid) and D (aspartic acid) are acidic amino acids.

**Supplementary figure 7.**
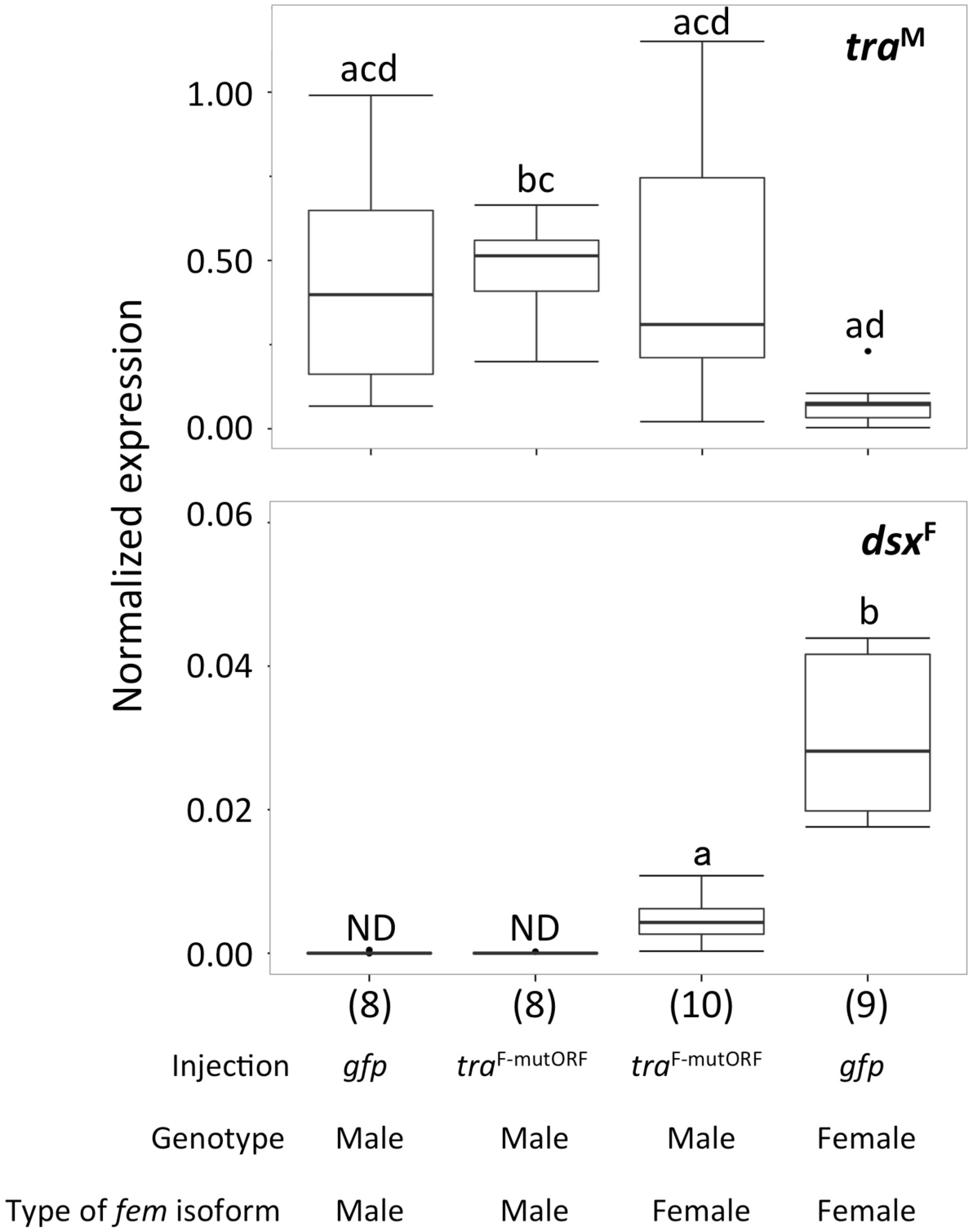
Normalized gene expression (median ± SD) of *tra*^M^and *dsx*^F^ inferred by qPCR. Individuals that grew to the first instar after injection of *tra*^F-mutORF^ and *gfp* mRNA at embryonic stage were used. According to treatments and results (Figure 8), males (individuals homozygous at both CSD loci) were classified into three groups: males treated with *gfp mRNA*, males without endogenous *tra*^F^expression after treatment of *tra*^F-mutORF^ mRNA and males with endogenous *tra*^F^expression after treatment of *tra*^F-mutORF^ mRNA. Females injected with *gfp* mRNA were used for comparison with these expression types between sexes. Primers for amplifying *tra*^M^ and *dsx*^F^ are shown in Figure 3A (white arrow) and in our previous study [37], respectively. Expression of *dsx*^M^ could not be detected at this developmental stage. Different letters indicate significant differences among groups (Tukey-Kramer test, P < 0.05). “ND” indicates “not detected”. Sample sizes are given in parentheses. Boxplots show the median and interquartile ranges (IQR) and 1.5 IQR.

**Supplementary figure 8.**
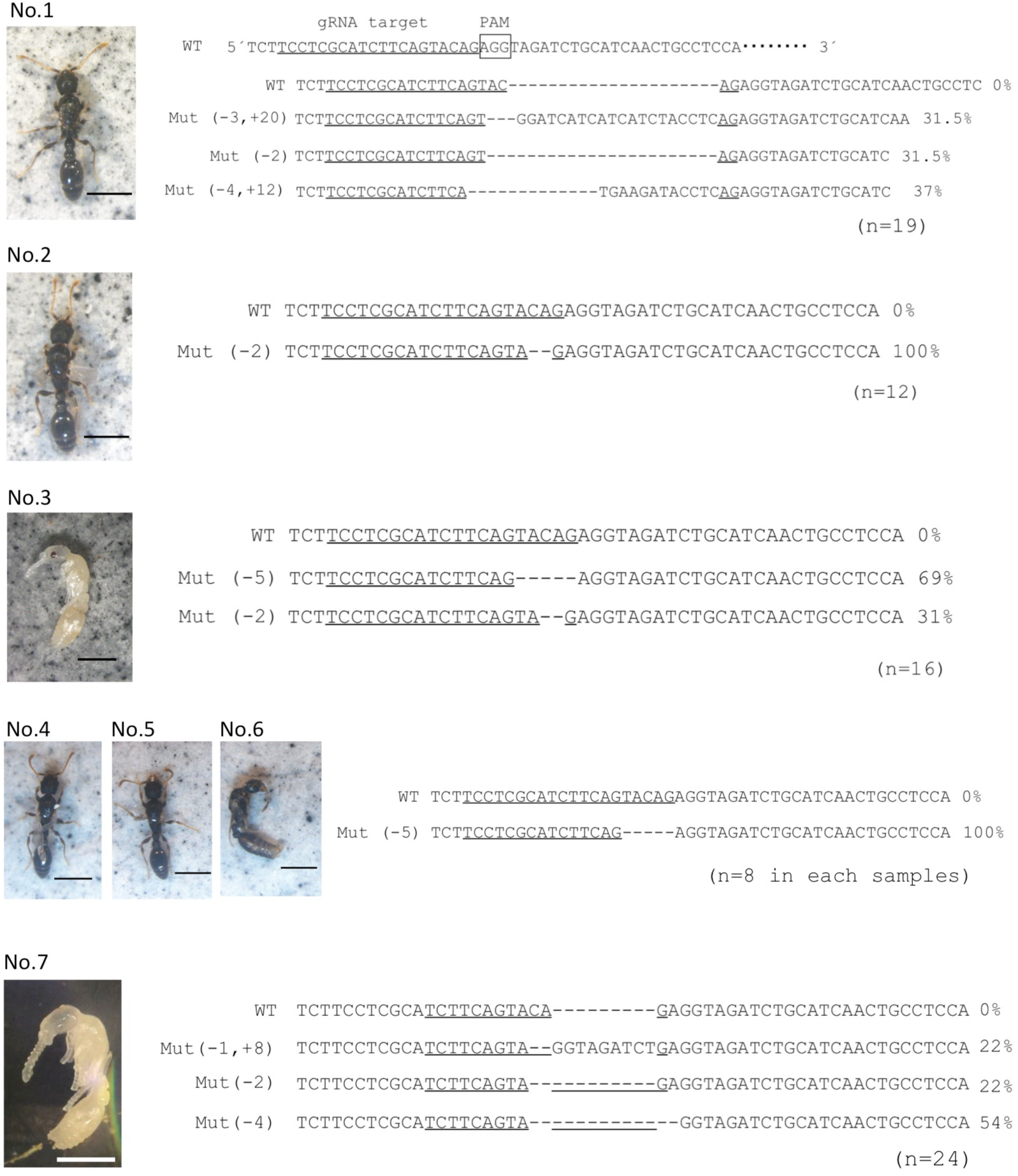
Detection of mutations in individuals that showed sex reversal from female to male after CRISPR/Cas9 mediation. Mutations detected in phenotypic males (n=7), heterozygous at two CSD loci. These individuals showed phenotypic characteristics of wild-type males (adults and pupae), but showed incomplete wing development. The scale bar indicates 1mm. The target site was in the third exon of the *tra* gene, which is located three nucleotides upstream of the PAM (AGG). The guide RNA (gRNA) targeting sequence is underlined. PCR products for the third exon of *tra* show that all seven phenotypic males had mutant alleles with frame shifts.

**Supplementary figure 9.**
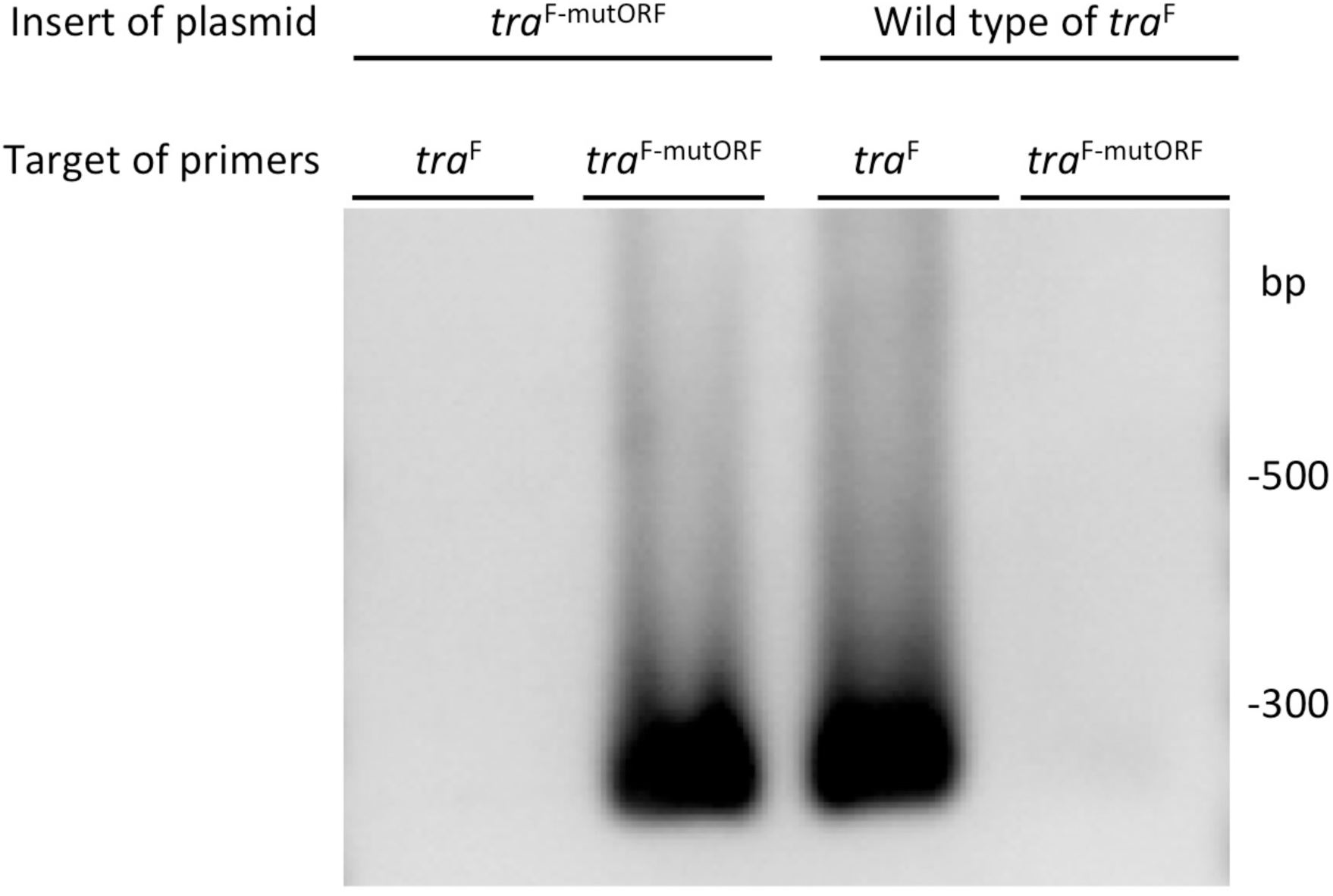
PCR to check primer specificity against endogenous *tra*^F^ mRNA. Primers designed to amplify endogenous *tra*^F^ mRNA (*Supplementary Table 1*) do not amplify plasmid template-inserted *tra*^F-mutORF^ mRNA, which introduced 12 synonymous single-nucleotide mutations. This primer set was used to check endogenous *tra*^F^ mRNA synthesis of males after injection of *tra*^F-mutORF^ mRNA.

**Supplementary Table 1.**
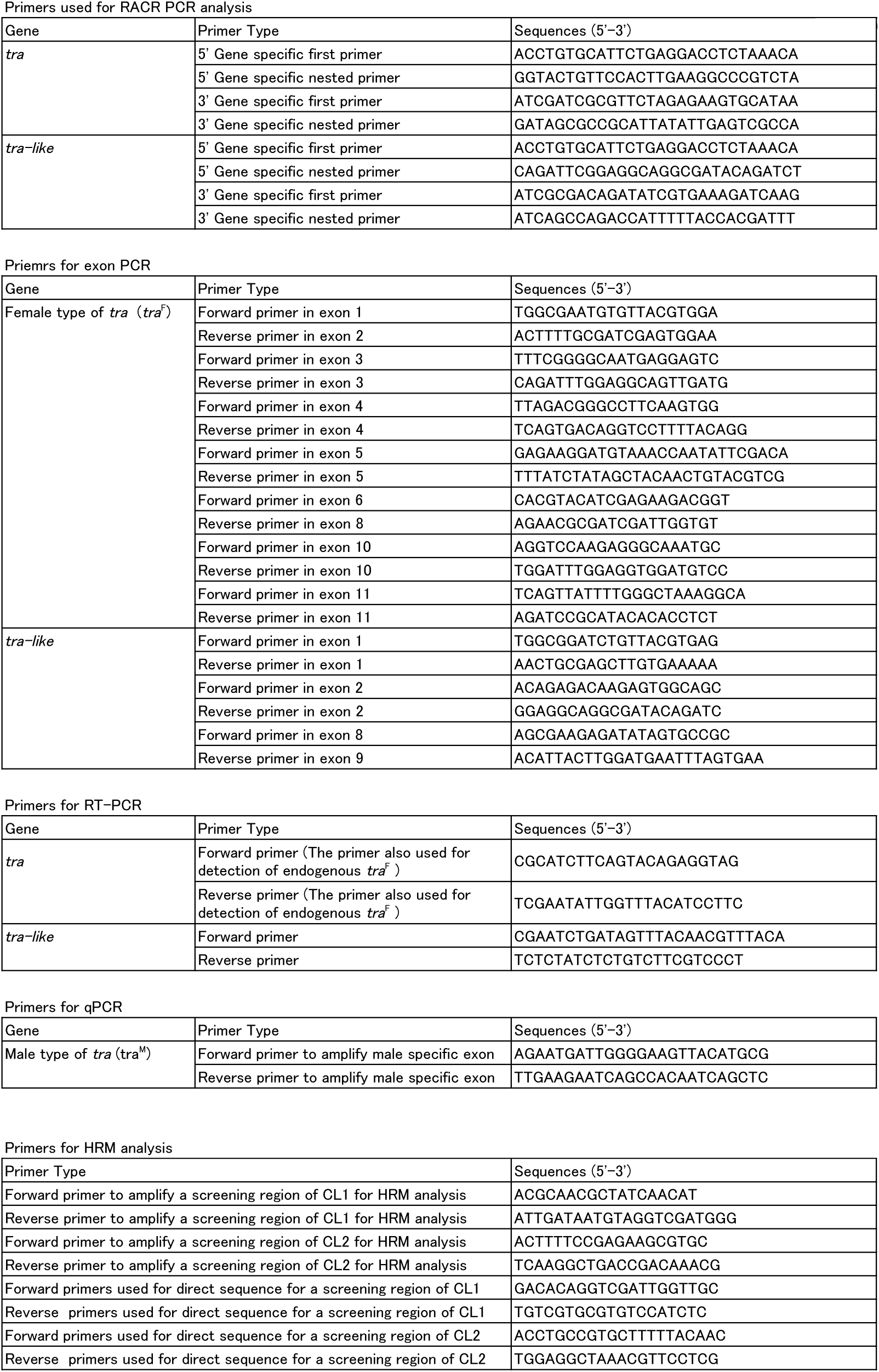
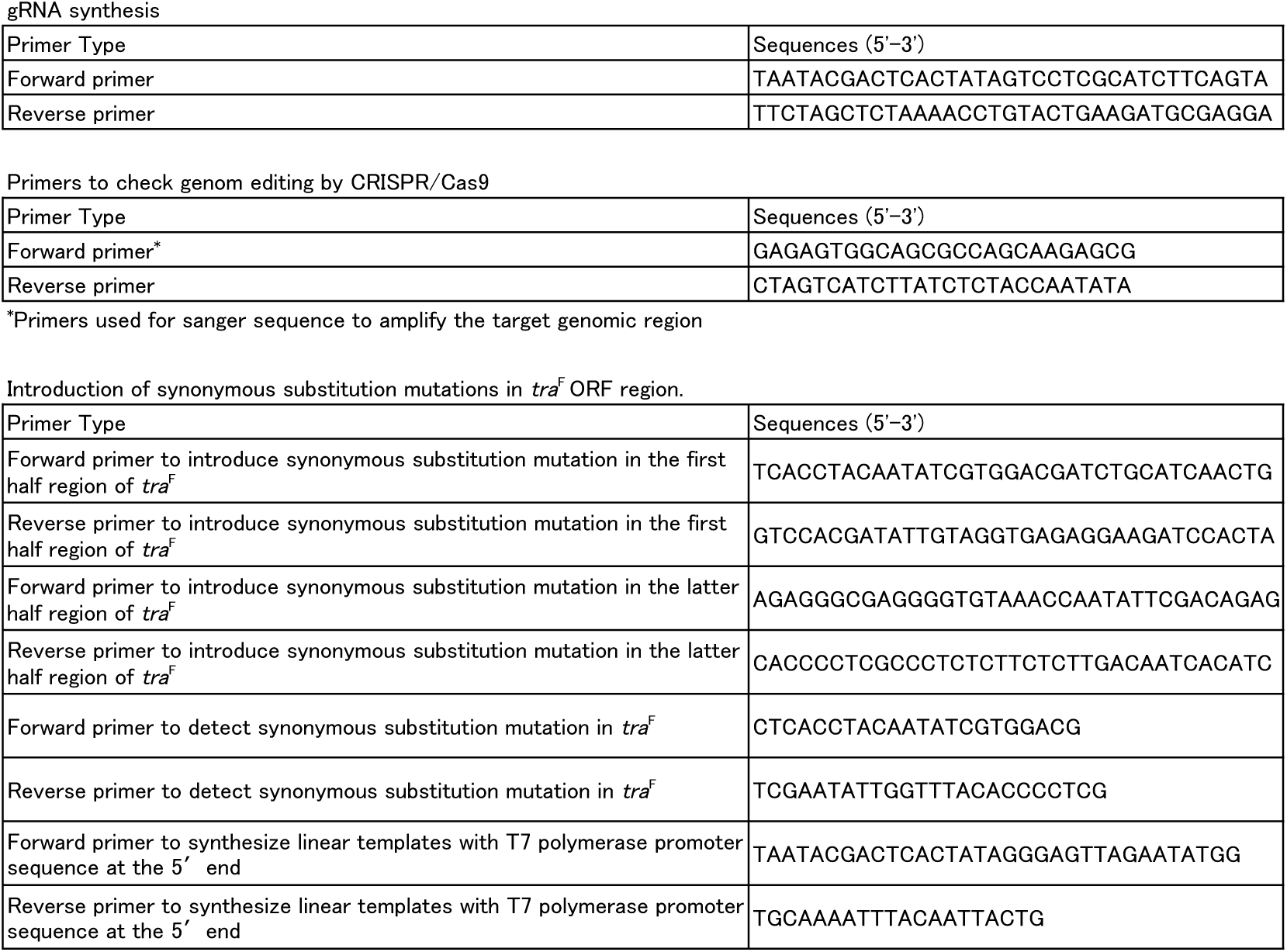
List of oligonucleotides used for RACE, RT-PCR, qPCR, and functional analysis.

**Supplementary Table 2.** Processing of endogenous *tra*^F^ in response to injection of *tra*^F-mutORF^ in early embryos (∼24 h).

| mRNA | Eggs injected | Eggs hatched | Genotype of CL1/CL2 |  |  |  | Ho/Ho specimens performed RT-PCR |  |  |
| --- | --- | --- | --- | --- | --- | --- | --- | --- | --- |
|  |  |  | He/He | He/Ho | Ho/He | Ho/Ho | Total | *tra*<sup>M</sup> isoform | *tra*<sup>F</sup> isoform |
| *tra*<sup>FmutORF</sup> | 608 | 55 | 14 | 12 | 11 | 18 | 18 | 8 (44%) | 10 (56%) |
| *gfp* | 583 | 77 | 17 | 24 | 16 | 20 | 19 | 19 (100%) | 0 (0%) |

**Supplementary Table 3.** Sex reversal (from female to male) of individuals treated with CRISPR/Cas9.

| RNPs |  | Injected eggs | Viability | Female | Male |
| --- | --- | --- | --- | --- | --- |
| sgRNAs<br>[ng/μl] | Cas9 protein<br>[ng/μl] | | | | |
| 500 | 600 | 716 | 4.5 % | 25 | 7 |

## Notes

### Competing Interest Statement

The authors have declared no competing interest.

